# CD2 expression acts as a quantitative checkpoint for immunological synapse structure and T-cell activation

**DOI:** 10.1101/589440

**Authors:** Philippos Demetriou, Enas Abu-Shah, Sarah McCuaig, Viveka Mayya, Salvatore Valvo, Kseniya Korobchevskaya, Matthias Friedrich, Elizabeth Mann, Lennard YW Lee, Thomas Starkey, Mikhail A. Kutuzov, Jehan Afrose, Anastasios Siokis, Michael Meyer-Hermann, David Depoil, Michael L. Dustin

## Abstract

The CD2 receptor has been described as an adhesion and costimulatory receptor on T cells. Here, transcriptional profiling of colorectal cancers (CRC) revealed a negative correlation between *CD2* expression and “exhausted CD8^+^ T-cells” gene signatures. Furthermore, we detected reduced surface CD2 levels in exhausted CD127^low^PD-1^hi^ CD3^+^CD8^+^ tumour infiltrating lymphocytes (TILs) in CRC. We describe a CD2 expression-level-dependent switch in CD2-CD58 localization between central and peripheral domains in the immunological synapse (IS). A peripheral “CD2 corolla” formed when CD2 surface expression was sufficiently high and its cytoplasmic domain intact. The corolla recruited other ligated receptors like CD28, boosted recruitment of activated Src-family kinases (pSrc), LAT and PLC-γ in the IS and consequently T-cell activation in response to a tumour antigen. Corolla formation and pSrc in the IS increased linearly with CD2 expression, whereas pSrc signals were reduced by high, “exhausted-like” levels of PD-1, which invaded the corolla. These results suggest two levels of inhibition of Src-family kinases in CD3^+^CD8^+^ TILs: reduced CD2 expression and high PD-1 expression.

## Introduction

Human CD2 (h)CD2 and its ligand CD58 were identified by monoclonal antibodies (mAbs) that inhibit T-cell conjugation with and killing of alloreactive target cells^1^. All mature human T-cells express CD2, with higher expression on memory T cells^2^. CD2 and CD58 interact through their extracellular immunoglobulin(Ig)-like domains, creating a complex of 13 nm length between cells^3^, similar to the TCR-pMHC complex. The cytoplasmic domain of CD2 is highly conserved and cross-linking human CD2 with mAbs activates T-cells in a manner that requires the intact CD2 cytoplasmic domain and TCR-CD3 complex^4^. In antigen-specific T-cell:antigen-presenting-cells (APC) conjugates CD2-CD58 interactions accumulate in the immunological synapse (IS) and contribute to sustained high intracellular calcium levels^5, 6^. In T-cells interacting with CD58-presenting supported lipid bilayers (SLBs), CD2-CD58 interactions form microdomains^7^. The microdomains are enriched in signalling molecules like phosphorylated lymphocyte-specific protein tyrosine kinase, (Lck) and Linker for activation of T cells (LAT) and excluded CD45 even in the absence of TCR engagement^7^. While there is a mouse homolog of CD2 (mCD2), there is no mouse CD58 and instead mCD2 binds to mouse (m)CD48. The CD2-CD58 interaction has a 50-fold higher affinity than mCD2-mCD48, and mCD2 does not interact with human CD58^8^. The CD2 contribution to T-cell activation has been associated with its adhesion^9^ and its signaling^4, 7, 10, 11, 12^ properties.

The CD2-CD58 pathway has been implicated in disease settings^13, 14, 15^. An allele of CD58 that decreases expression has been associated with increased susceptibility to multiple sclerosis^16^ and *CD2/CD58* genetic variants have been associated with rheumatoid arthritis risk^17^. It has been shown that during *in vitro* chronic activation of human CD8^+^ T-cells, CD2 ligation prevents development of an exhausted CD127^low^PD1^hi^ phenotype^15^. The related CD2 transcriptional signature has been associated with better viral clearance, but more severe autoimmunity^15^. In the context of cancer, loss of CD58 expression has been reported mainly in blood malignancies^18^. Two studies have reported that blockade or loss of CD2-CD58 interactions reduced the *in vitro* cytotoxicity of human CD8^+^ T cells to melanoma cell lines^13, 14^. To our knowledge, there have been no reports on CD2 levels in tumour infiltrating lymphocytes (TILs). Similarly, the interplay between CD2-CD58 interaction and Programmed cell death protein-1 (PD-1)-Programmed death-ligand-1 (PD-L1) interaction, which has been implicated in attenuated responses from exhausted TILs ^19, 20, 21, 22, 23^ is unknown.

We hypothesized that exhausted TILs have reduced CD2 levels and this contributes to their attenuated anti-tumour responses. We tested our hypothesis by determining the phenotype of CD8^+^ TILs from colorectal cancer (CRC) patients. CD8^+^ TILs from most CRC patients tested have a lower number of CD2 molecule per cell compared to antigen-experienced T-cells from peripheral blood of healthy individuals. This prompted us to ask how do CD2 levels affect its function and subsequently T-cell responses. We, thus, investigated the properties of the CD2-CD58 interaction upon antigen recognition, synapse formation and T-cell activation. Our study revealed an unexpected and unique contribution of CD2 levels to TCR signalling and protein organisation at the IS and thus T-cell activation status.

## Results

### CD8^+^ TILs and in particular exhausted TILs, from CRC patients, express low levels of CD2

Colorectal cancer is the third most common type of cancer worldwide with a high mortality rate. Interrogation of transcriptomic profiling data from CRC patients (n=255) in The Cancer Genome Atlas (TCGA) identified that the expression of CD8^+^ T-cell exhaustion signatures negatively correlated with *CD2* expression. This was true for both genesets in “Exhausted vs Effector CD8 T-cell”, (GSE 41867^24, 25, 26^, *p<*0.001), (Figure 1a, top) and “Exhausted vs. Naïve CD8 T-cell” (GSE 41867, *p<*0.001), (Figure 1a, bottom). This finding was then validated using an independent CRC cohort (METALLIC) of 148 patients with stage III CRCs with whole transcriptomic profiles. Linear regression confirmed a similar negative correlation between CD8^+^ T-cell exhaustion signatures and *CD2* in this cohort (p<0.001 for both, Supplementary Figure 1). Further analysis in TCGA revealed that this negative correlation was independent of microsatellite instability (MSI), which correlates with CD8^+^ T-cell infiltration in CRC^27^.

**Figure 1.**
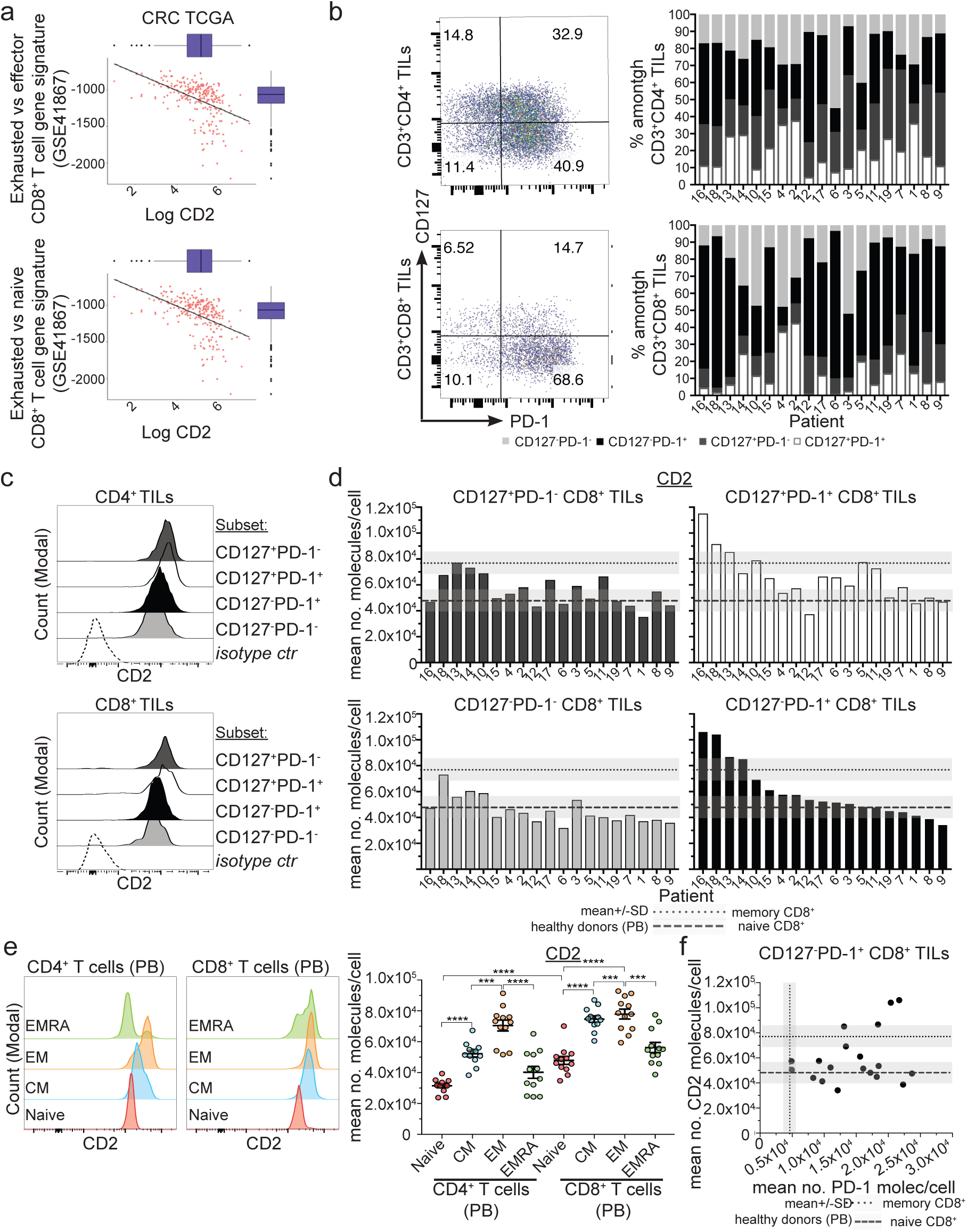
CD8^+^ TILs from CRC patients, including exhausted CD127^low^PD-1^hi^ CD8+ TILs, can show a CD2^low^ phenotype. **a)** The correlation between exhausted CD8^+^ T-cell gene signature and CD2 expression from whole transcriptomic analysis of CRC patients from The Cancer Genome Atlas (TCGA) is shown (n=255) for (top) “exhausted vs effector CD8^+^ T cell gene signature” and (bottom) “exhausted vs naive CD8^+^ T cell gene signature”. **b)** The proportions of CD127^+^PD-1^−^, CD127^+^PD-1^+^, CD127^−^PD-1^+^, CD127^−^PD-1^−^ cells present within the viable CD3^+^CD4^+^ (top) and CD3^+^CD8^+^ gates (bottom) of CRC TILs are shown. Representative CD127, PD-1 expression profiles are shown (left) and the ratio of each subset for each one of 19 CRC patients (right). **c)** Representative histograms of CD2 profiles of (top) CD4^+^ and (bottom) CD8^+^ TILs from (b) are shown. **d**) The mean number of CD2 molecules/cell, for the CD8^+^ TIL subset shown in (c), is shown for each patient. Patients were sorted from highest to lowest CD2 expression of the CD127^−^PD-1^+^ T-cell compartment. **e)** (Left) Representative CD2 expression profiles, from 1 of 12 healthy donors, for CD4^+^ and CD8 T-cell subsets based on their CD62L and CD45RA; Naïve (CD62L^+^CD45RA^+^), central memory, CM (CD62L^+^CD45RA^−^), effector memory, EM (CD62L^−^CD45RA^−^) and effector memory that re-express CD45RA (EMRA) (CD62L^−^CD45RA^+^). (Right) The mean CD2 expression levels (molecules/cell) of 12 healthy donors and of each individual donor are shown. **, p value <0.007, ***, p <0.007 ****, p value <0.0001, with parametric Welch’s *U-*test. **f)** CD2 against PD-1 levels of CD127^−^PD-1^+^ CD8^+^ TILs from (d) are shown for each donor. Dotted line crossing x-axis (with grey rectangle) represents the average PD-1 levels (±SD) expressed in memory PD-1^+^ CD3^+^CD8^+^ T cells found in peripheral blood of healthy controls. Dashed or dotted line crossing y-axis (with grey rectangle) represents the average CD2 levels (±SD) expressed in naive and memory CD3^+^CD8^+^ T cells, respectively, found in peripheral blood of healthy individuals.

The transcriptomic findings were corroborated by analysing single cell suspensions obtained from primary CRC tissue resections and profiled using flow cytometry. We determined the absolute levels of surface CD2 expression in CD4^+^ and CD8^+^ TILs subdivided based on their CD127 and PD-1 expression profile (Figure 1b-d and Supplementary Figure 1) from 19 CRC patients (Supplementary Table 1). In 13 out of 19 CRC patients, >50% of CD8^+^ TILs were CD127^−^PD-1^+^ whereas only 1 out of 19 CRC patients showed such composition among CD4^+^ T-cells (Figure 1b).

CD2 levels in TILs were next compared to that of other T-cell populations. The absolute number of CD2 molecules on naïve, central memory, effector memory and effector memory that re-express CD45RA (EMRA) from peripheral blood of 12 healthy individuals was determined (Figure 1e). We found that naïve CD4^+^ T-cells express the lowest number of CD2 molecules/cell (3.2×10^4^±4.4×10^3^), whereas effector memory CD8^+^ T-cells express the highest number of CD2 molecules/cell (7.8×10^4^±1.1×10^4^), a 2.4-fold range (Supplementary Table 2). This data is in line with a previous report showing that human memory CD4^+^ or CD8^+^ T-cells express higher levels of CD2 than their naïve counterparts^2^. CD2 levels in TILs were 3-fold more variable (s.d./mean) compared to their counterparts in healthy individuals (Supplementary Table 3). A substantial number of CRC patients had CD8^+^ TILs with CD2 levels that were within and/or lower than the number of CD2 molecules/cell on peripheral blood naïve CD8^+^ T-cells from healthy individuals; particularly in CD127^+^PD-1^−^, CD127^−^PD-1^+^ and CD127^−^PD-1^−^ subsets. The lower number of CD2 molecules/cell will be referred to as “CD2^low^ phenotype”. Focusing on CD127^−^PD-1^+^ CD8^+^ TILs, 14 out of 19 CRC patients exhibited the CD2^low^ phenotype (4.8×10^4^±7.6×10^>^; Group A;) which was significantly different (p<0.0001) than CD2 levels in memory CD8+ T cells (7.6×10^4^±8.9×10^3^) from peripheral blood of healthy individuals (Supplementary Figure 1). CD127^−^PD-1^+^ CD8^+^ TILs in the remaining 5 CRC patients (Group B) expressed 1.8-fold higher CD2 levels (9.0×10^4^±1.5×10^4^) similar to the later memory CD8^+^ T cells. This smaller number CD2 molecules/cell is not due to the presence of naïve CD8^+^ T-cells as two of these subsets also express PD-1, an inhibitory receptor expressed on antigen-experienced T-cells^28^. Furthermore, >95% of TILs expressed the memory T-cell marker CD45RO.

To determine whether the CD127^−^PD-1^+^ T-cell compartment consisted of exhausted CD127^low^PD-1^hi^ T-cells, we performed absolute quantification of PD-1 of this subset and compared this to PD-1 levels in PD-1^+^ memory T-cells found in peripheral blood of healthy individuals (Figure 1f and Supplementary Table 4-5). This revealed that in 17 out of 19 CRC patients, CD127^−^PD-1^+^ CD8^+^ TILs had a higher number of PD-1 molecules/cell compared with the mean levels of PD-1 (5.9×10^3^±1.2×10^3^ molecules/cell) in peripheral blood of healthy individuals (mean number of PD-1 in 19 CRC patients was 1.45×10^4^±5.58×10^3^ molecules/cell). In fact, in 12 out of these 17 patients their CD127^low^PD-1^hi^ CD8^+^ TILs exhibited the CD2^low^ phenotype. This observation supports the results from the CRC whole transcriptomic analysis (Figure 1a) that identified a negative correlation between CD8^+^ T-cell exhaustion gene signatures and *CD2* expression. Interestingly, this was the opposite case for CD127^−^PD-1^+^ CD4^+^ TILs. In 17 out of 19 CRC patients, CD127^−^PD-1^+^ CD4^+^ TILs had indeed high PD-1 levels yet their CD2 expression was within or higher than the range of CD2 molecules/cell in peripheral blood memory CD4^+^ T-cells from healthy individuals (Supplementary Figure 1 and Supplementary Table 4-5). The CD2^low^ phenotype was detected mainly in CD127^−^PD-1^−^ CD4^+^ TILs. The impact of low CD2 levels on exhausted TILs is unclear. These observations motivated us to undertake a more thorough investigation of the basic biology of CD2 in T-cells and the functional role of CD2 expression. Therefore, we moved forward to investigate how CD2 numbers affect synapse formation, TCR signalling and T-cell activation.

### A unique ring pattern, “corolla”, formed by CD2-CD58 interactions in the IS

Firstly, we set out to determine the localization of CD2 when a mature T-cell synapse forms in a cell conjugate model system. Human CD4^+^ and CD8^+^ T-cells, isolated from PBMCs, were transfected with *6F9* (MHC-II restricted) or *1G4* (MHC-I restricted) TCR, as described in^29^ and conjugated to MAGE-A3_243-258_ or NY-ESO-9V_157-165_ loaded CF996 EBV-transformed B-cells. 72% of cell conjugates imaged had a CD2 signal and this showed an accumulation in the interface between the T-cell and the B-cell. CD2 and CD11a, the α-subunit of LFA-1, staining revealed the presence of a CD2 signal in the distal regions of the cell-cell interface, outside the LFA-1-enriched peripheral supramolecular activation cluster (pSMAC) or within the pSMAC with segregation of LFA-1 and CD2 signals (Figure 2a-b and Supplementary Figure 2). These 3D images argued for an unexpected localization of CD2 in the IS, distinct from previous description of cSMAC localization^30, 31^.

**Figure 2.**
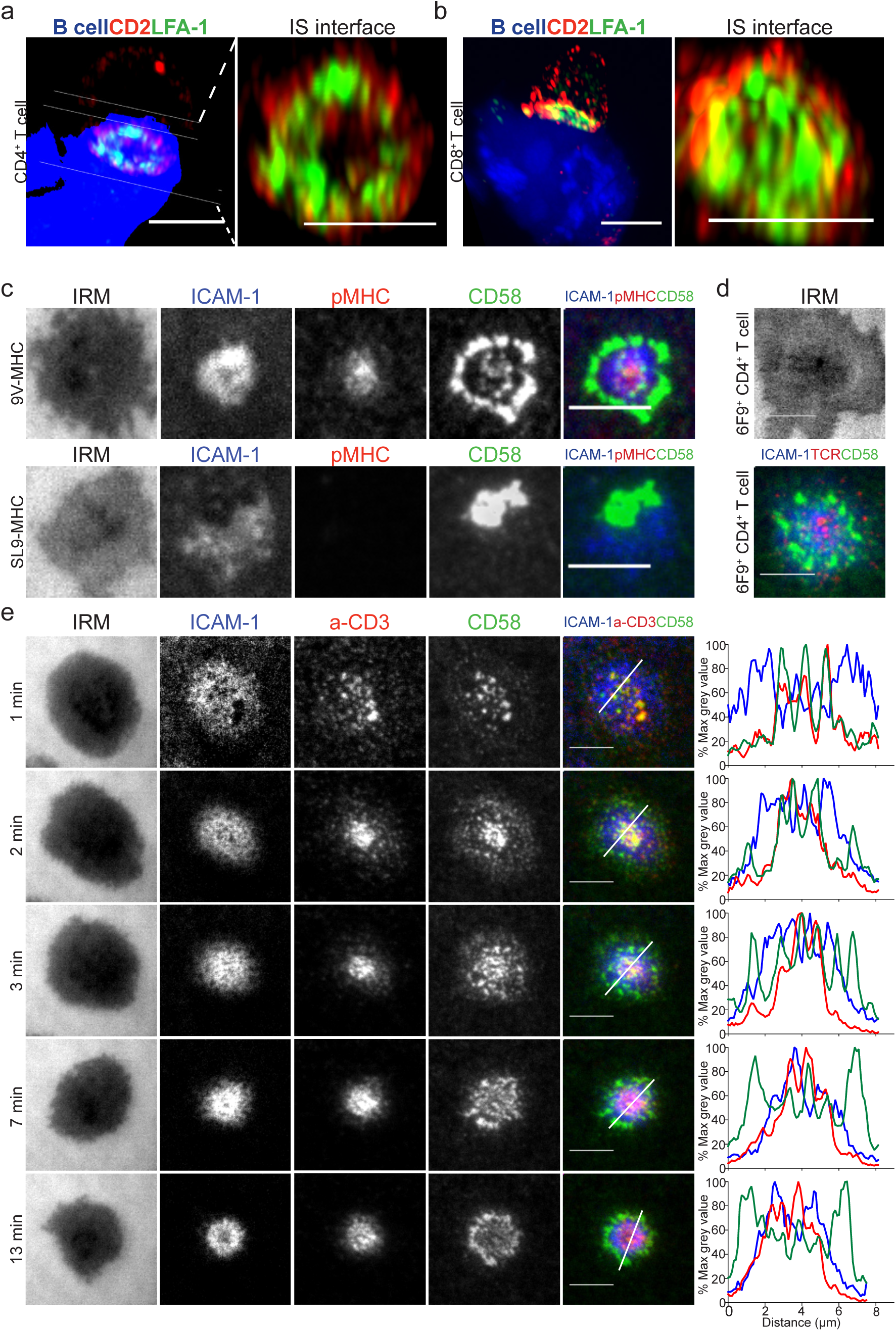
A unique ring pattern, “corolla”, formed by CD2-CD58 interactions in the IS. **a-b)** 3D rendering images (IMARIS software) of T:B cell conjugates and enlarged 1 μm thick slice (white solid lines) of the T:B cell IS interface. *6F9*^+^ CD4^+^ T-cells (a) and *1G4*^+^ CD8^+^ T-cells (b), were conjugated with EBV-transformed B cells (blue-volumetric dye) pulsed with 1 μM MAGE-3A_243-258_ or 1μM NY-ESO-9V_157-165_ peptide, respectively, for 25-30 min. CD2 (red) and LFA-1-α-subunit (green) staining shown. Images were captured on an Airyscan confocal microscope (ZEISS). Representative images are shown from two independent experiments. **c)** 1G4^+^ CD8^+^ T-cells, fixed 15 min post-incubation on ICAM1 (200/μm^2^), CD58 (200/μm^2^) reconstituted SLBs, in addition to (top panel) NY-ESO-9V-peptide-loaded HLA-A2 (30/μm^2^) (top panel) or (bottom panel) GAG-SL9-peptide-loaded HLA-A2 (30/μm^2^). Cells were imaged with TIRFM. **d)** *6F9*^+^ CD4^+^ T-cells, treated same as in (c) but on MAGE-A3 peptide loaded HLA-DP4 (30/μm^2^) instead. *6F9*^+^ TCR CD4^+^ T-cells were stained with a fluorescently labeled anti-mouse TCRβ Fab (H57 clone) prior to SLB incubation. **e)** CD4^+^ T-cells, incubated on ICAM1 (200/μm^2^), anti-CD3 Fab (30/μm^2^), CD58 (200/μm^2^) reconstituted SLBs. Cells were imaged at 4s intervals with TIRFM. Representative images within 13 min from initial contact are shown. Histograms depict the intensity profiles on the diagonal white lines in overlay images. Raw pixel intensity signal normalized to maximum intensity pixel of each channel. A representative experiment of three independent experiments with either CD4^+^ or CD8^+^ T-cells shown. IRM, interference reflection microscopy, signal shows the spreading phase of the cell. Scale bar, 5 μm.

This was an unexpected organization as one would have expected the CD2-CD58 to be present in the same area as the TCR-pMHC^32^. To gain a deeper understanding of this puzzling localization we turned to the use of the supported lipid bilayer (SLB)^33, 34^ reconstituted with pMHC, ICAM1 and CD58. The number of CD58 molecules on the CF996 B-cell line, primary human monocytes and monocyte-derived dendritic cells was determined (Supplementary Figure 2 and Supplementary Table 6); and based on this, the density of CD58 used on SLBs, was 200 molecules/μm^2^. SLBs presenting distinctly fluorescently labelled ICAM1 (200/µm^2^) and CD58 (200/µm^2^) with monomeric pMHC (30/µm^2^) were used to stimulate antigen specific T-cells, which were then fixed at 15 min (the timepoint typically used to study mature IS on SLBs similar to the cell conjugate assay) and imaged by total internal reflection fluorescence microscopy (TIRFM). *1G4* TCR transfected CD8^+^ T-cells formed IS with SLB containing NY-ESO-9V-peptide loaded HLA-A2, ICAM1 and CD58; similarly, 6*F9* TCR transfected CD4^+^ T-cells formed synapses on SLB with MAGE-A3-peptide loaded HLA-DP4 SLBs (Figure 2c-d). We found that CD2-CD58 interactions were localized distal to the LFA-1-ICAM1 pSMAC in an array of flower petal-like microdomains that collectively we define as a “corolla”; closely resembling the CD2 distribution in the cell conjugates. *1G4* TCR transfected CD8^+^ T-cells didn’t form this corolla pattern on SLB presenting the non-cognate GAG-SL9-peptide loaded HLA-A2. In this setting, the CD2-CD58 interactions tended to accumulate in a single microdomain (Figure 2c). This antigen-independent CD2-CD58 interaction excluded LFA-1-ICAM1 interactions similar to the CD2 corolla, but localized to one side of the LFA-1-ICAM1 focal zone rather than surrounding an LFA-1-ICAM1 enriched pSMAC. These data indicate that productive TCR engagement is required for CD2 corolla formation. Thus, we identified a unique pattern of CD2 in the IS between T and B cells and reconstituted the pattern in SLB presenting only pMHC, ICAM1 and CD58.

### Dynamics of corolla formation

The use of anti-CD3 Fab to engage TCR enabled the use of untouched polyclonal CD4^+^ T-cells isolated from human peripheral blood. Imaging of the SLB, also containing fluorescently labelled ICAM1 (200/µm^2^) and CD58 (200/µm^2^), was initiated as the T-cells settled and made first contact. T-cell spreading was complete within 1 min and engaged TCR-anti-CD3 and CD2-CD58 interactions were initially co-localized in microclusters (Figure 2e and Supplementary Video 1-2). As soon as spreading was complete the TCR/CD2 microclusters moved towards the center of the IS as reflected at the 2 min time point, with some domains enriched in CD2 only beginning to appear. At 3 min of synapse formation, CD2-CD58 microdomains re-emerged in the LFA-1-ICAM1-enriched pSMAC and outside of the pSMAC i.e. in the dSMAC, but there was still evidence of TCR-anti-CD3 interactions within intensifying CD2-CD58 microclusters. At 7 min a new pattern emerged as CD2-CD58 interactions were reduced in the TCR-enriched cSMAC. The CD2-CD58 microdomains that emerged in the periphery lacked correlated TCR-anti-CD3 accumulation and the distinct flower petal-like “corolla” formed around the pSMAC. The corolla pattern, then, persisted through 13 min. The CD2-CD58 corolla petals were visibly segregated from LFA-1-ICAM1 interactions. New TCR-anti-CD3 microclusters appear to emerge from the corolla petals (Supplemental movies 1-3) suggesting that CD2-CD58 interactions have an ongoing role in new TCR-anti-CD3 interactions. Movement of CD2-CD58 interactions was biphasic with the initial phase showing strong co-localization and tracking with TCR followed by expansion into the pSMAC and dSMAC, the region associated with TCR signal initiation^35^. Finally, these CD2-CD58 dynamics are very different from other costimulatory receptors like CD28 and PD-1, which have been described to localize to the cSMAC^20, 21, 36, 37^. How do these other costimulatory-inhibitory receptors behave in the presence of the CD2 corolla?

### The corolla captures ligated CD28 and Inducible T-cell Costimulator (ICOS)

The signaling and biochemical pathways triggered by CD28 have been compared to CD2 induced pathways either when both receptors are engaged together or separately^12, 22, 23, 38^. However, there have been no studies looking at the localization of CD28 and CD2 when both are ligated by freely diffusing CD58 and CD80 during IS formation. This becomes even more interesting in light of our identification of the CD2 corolla. CD28 engagement by CD80 in the context of ICAM1 and agonist pMHC only generates peripheral CD28-CD80 microclusters in the first min of IS formation that coalesce into a tight ring around the synaptic cleft at later stages (20 min), which make up the cSMAC^37, 39^. In our study, in order to track ligated CD28 in the presence and absence of CD2 corolla formation, a fluorescently-tagged CD80 was added to the SLB presenting ligands.

In the absence of CD58, 98.4(±1.1)% CD4^+^ T-cells displayed an annular ring of CD28-CD80 interactions within the cSMAC (Figure 3a, >400 cells from 4 donors), as previously described^37, 39^. In the presence of CD58, CD28-CD80 interactions were recruited to the corolla in 90.8(±6.6)% of n=4 CD4^+^ T-cells (Figure 3a and Supplementary Movie 3). Similar results were obtained with CD8^+^ T-cells (99.7(±0.4)% corolla positive, n=4 donors). This phenotype was reproduced in an antigen-specific SLB system, using *1G4* TCR transfected CD8^+^ T-cells (97.9% corolla positive, n=2 donors). (Supplementary Figure 3). Similarly, ICOS-ICOS-L interactions were localized throughout the cSMAC in the absence of CD58 and in CD2 corolla in the presence of CD58 in the SLB (Figure 3b). Therefore, CD2-CD58 interactions relocate classical costimulatory receptor-ligand pairs to the corolla.

**Figure 3.**
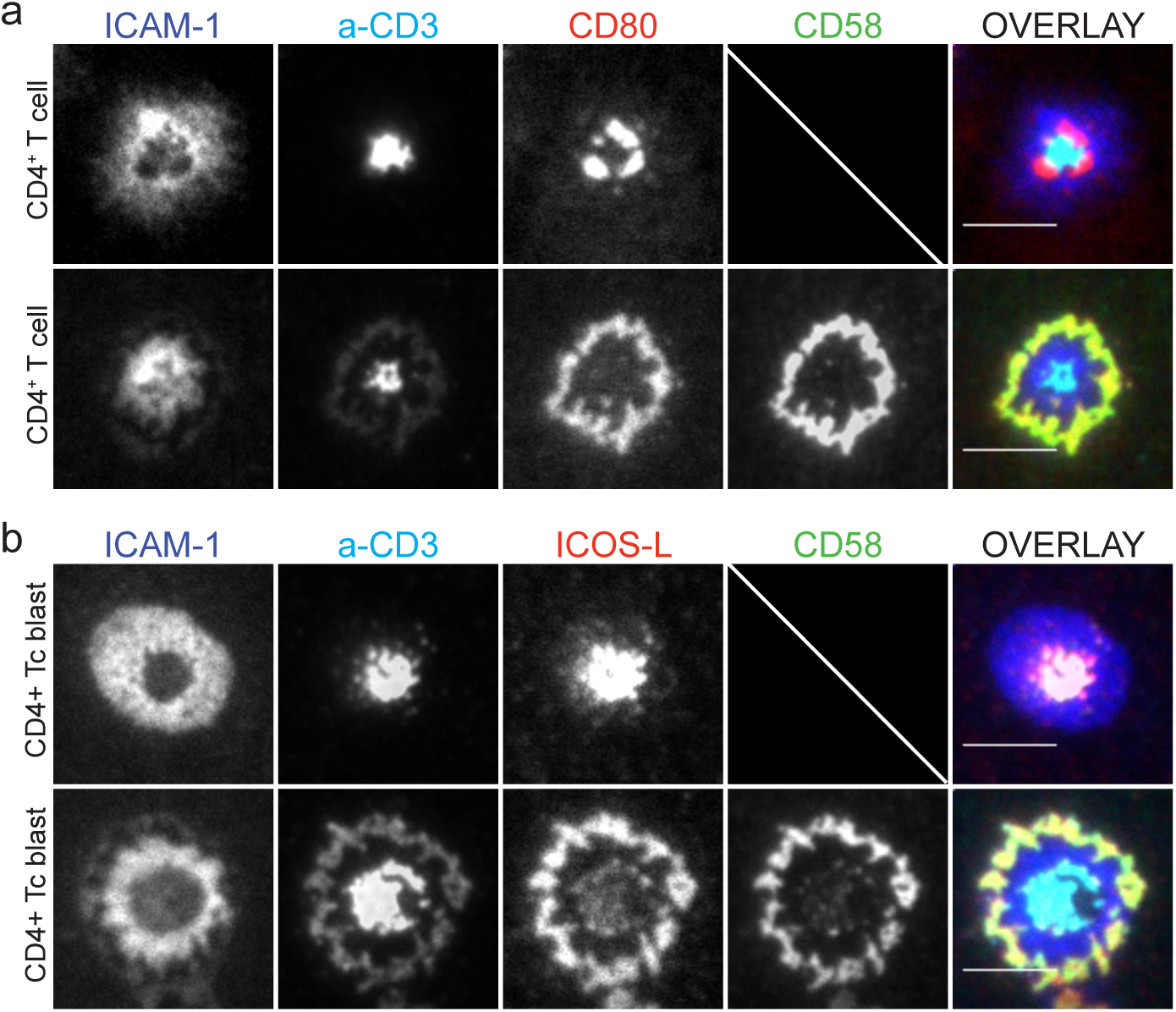
The corolla captures ligated CD28. **a)** Human CD4^+^ T-cells incubated on ICAM1 (200/μm^2^), anti-CD3 Fab (30/μm^2^), CD80 (100/μm^2^) without (top) or with CD58 (200/μm^2^) (bottom) reconstituted SLBs and fixed at 15 min. **b)** Activated human CD4+ T cell blasts incubated on ICAM-1 (200/μm^2^), anti-CD3 Fab’ (30/μm^2^), ICOS-L (100/μm^2^) without (top) or with CD58 (200/μm^2^), (bottom) reconstituted SLBs and fixed 15 min. Cells were imaged with TIRFM and representative images are shown from four independent experiments. Scale bar, 5 μm.

### CD2 expression levels determine corolla formation

Given the unexpected localization of CD2-CD58 interactions and the spatial effect of CD2 corolla on other costimulatory receptors we sought to determine the requirements for corolla formation other than the presence of TCR agonists and CD58. Due to differential CD2 expression levels in different T-cell populations (Figure 1), we hypothesized that the number of CD2 molecules on the cell surface controls corolla formation. We exploited the four major natural T-cell subsets, naïve (CD62L^+^, CD45RA^+^), and memory (CD45RA^−^) for CD4^+^ and CD8^+^ T-cell that varied in their levels of surface CD2 (Supplementary Table 2) and asked how corolla formation varies in these (Figure 4a-b). Consistent with lower CD2 numbers in naïve subsets, we detected lower levels of accumulated CD58 in the IS formed by naïve subsets compared to their memory counterparts (Figure 4a-c). Naïve CD4^+^ T-cells, with the lowest number of CD2, had the least intense CD2 corolla with smaller clusters sparsely arranged around the pSMAC (Figure 4a). In fact, many of the naïve CD4^+^ T-cells did not arrange CD2 clusters in a corolla at all. Instead, CD2 clusters were mainly within the cSMAC and the inner border of the LFA-1-ICAM1-enriched pSMAC; hence we detected only 60(±4)% (SEM) corolla positive cells in this subset (see Figure 4 legend how ‘corolla’ is scored). Naïve CD8^+^ T-cells were 83(±4)% corolla positive while memory CD4^+^ T-cells and memory CD8^+^ T-cells, 94%(±4)% and 90(±4)%, respectively (Figure 4d). Memory CD4^+^ or CD8^+^ T-cells accumulated larger and brighter CD2 microdomains, with many of them having contiguous band of CD2-CD58 interactions around the pSMAC compared with their naive counterparts.

**Figure 4.**
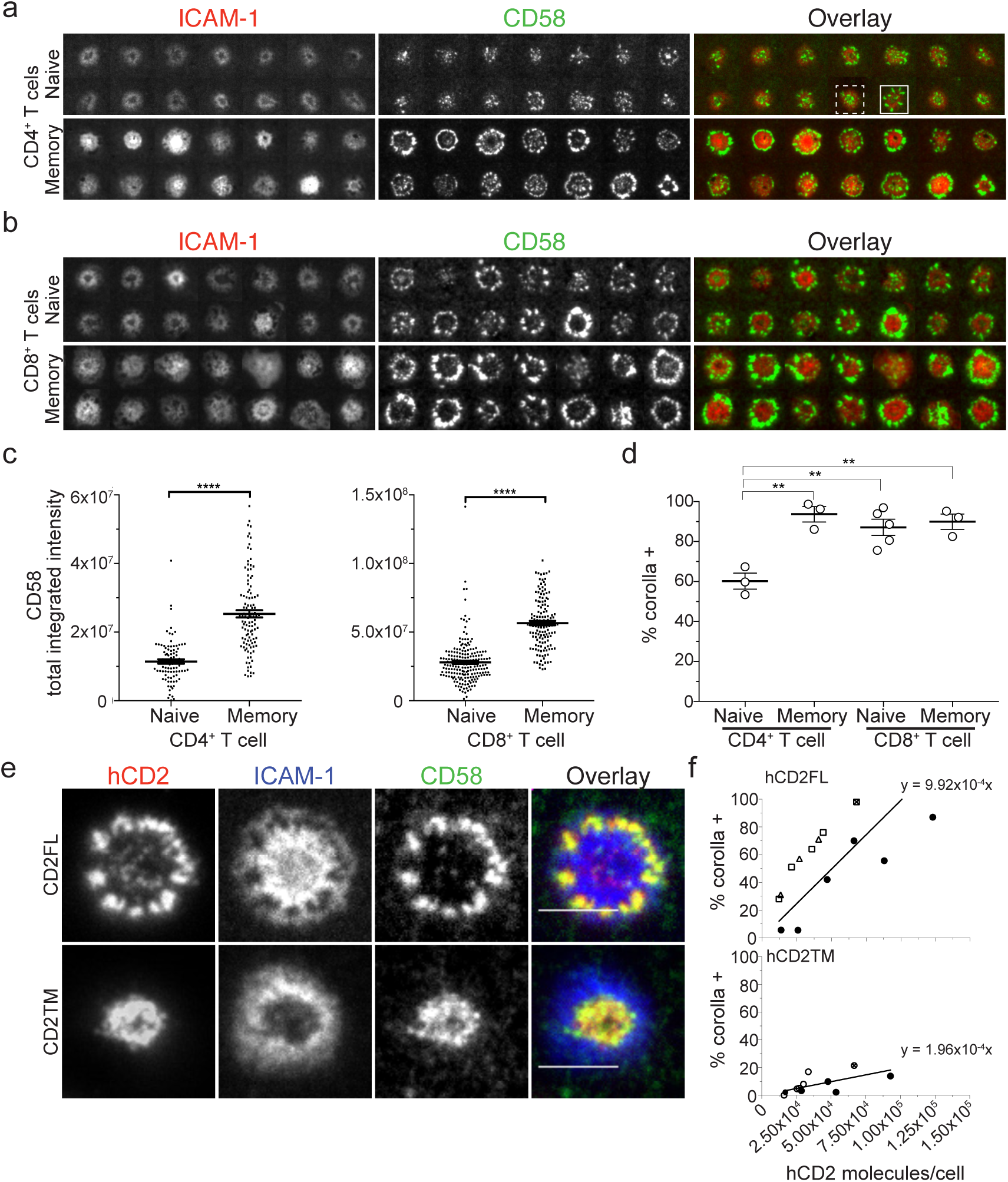
CD2 expression determines corolla formation. **a)** A random image selection of naïve (CD62L^+^CD45RA^−^) or memory (CD45RA^−^) CD4^+^ T-cells incubated on ICAM1 (200/μm^2^), anti-CD3 Fab (30/μm^2^) and CD58 (200/μm^2^) reconstituted SLBs, fixed at 15 min and imaged with TIRFM; examples of corolla positive (solid line rectangle) and corolla negative (dashed line rectangle) are shown. **b)** Same as in (a) but with CD8^+^ T-cells. **c)** Quantification (from experiments as in (a)) of total CD58 accumulation in the IS of CD4^+^ and CD8^+^ T-cell subsets. ****, p value <0.0001 with non-parametric Mann Whitney *U-*test. A representative of three experiments is shown. Error bars represent standard error of mean, SEM. **d)** Percentage of T-cells forming an IS with a CD2 corolla (corolla^+^) in experiments as in (c-d), was determined for naïve and memory T-cell subsets. Circles represent donors. **, p value <0.006, with parametric Welch’s *U-*test. 80-200 cells/subset were used for quantification. Error bars represent SEM. **e)** Representative images of AND T-cells expressing either human CD2 full length (hCD2FL) or hCD2 lacking the cytoplasmic tail (hCD2TM), at levels comparable with CD2 levels found in human peripheral blood T-cells, on ICAM1 (200/μm^2^), MCC-I-E^k^ (30/μm^2^) and CD58 (200/μm^2^) reconstituted SLBs, fixed 15 min post-incubation and imaged with TIRFM. **f)** The percentage of hCD2FL- or hCD2TM-expressing AND T-cells that formed a CD2 corolla (% corolla^+^) in experiments such as in (e) over a range of hCD2FL (top) and hCD2TM expression (bottom). At least 100 cells were considered/data point. 95% CI of regression lines for FL, 7.15×10^−4^ to 12.69×10^−4^; for TM 1.17×10^−4^ to 2.75×10^−4^. Each different symbol represents one data point from an individual experiment.

Corolla formation indeed appears to correlate with CD2 numbers in the natural subsets, however, there are other differences between the subsets that could account for the differences in the IS structure. This led us to develop a system in which transient expression of different average numbers of CD2 in a single T-cell type could be used to determine the relationship between the levels of CD2 and corolla formation. Mouse AND T-cells were used as a T-cell host lacking human CD2 (hCD2) activity. Varying levels of two versions of hCD2 were expressed: the full length with the complete cytoplasmic domain of 116 residue (hCD2FL) or a truncated form of hCD2 with only a 6-residue cytoplasmic domain (hCD2TM). The titration of hCD2 expression was achieved by electroporating varying amounts of mRNA encoding hCD2FL or hCD2TM in the AND T-cells such that the range of hCD2 molecules/cell also spans the ranges we observed in human T-cells. The mean number of hCD2 molecules in electroporated AND T-cells was quantified by flow cytometry (Supplementary Figure 4). AND T-cells expressing 1.2×10^4^-4.4×10^4^ molecules/cell of hCD2FL were incubated on ICAM1, pMHC and CD58 reconstituted SLBs and corolla formation was scored at 15 min. The percentage of cells forming a corolla increased with increasing levels of hCD2FL (Figure 4e-f and Supplementary Figure 4). hCD2TM also showed a linear relationship between corolla formation and CD2 number per cell but was significantly less efficient (5.1-fold) in forming a CD2 corolla compared with hCD2FL (Figure 4e-f and Supplementary Figure 4). These results establish that the amount of CD2 expressed and the cytoplasmic domain are both determinants of corolla formation. Thus, TILs with a CD2^low^ phenotype are likely to have less CD2-CD58 interactions during IS formation and corolla formation.

### CD2 corolla amplifies TCR and LAT dependent signals and enhances T-cell activation

We have described a role for CD2 as a master organiser of costimulatory receptors, CD28 and ICOS, in the IS. We sought to investigate the implications of CD2 organisation on its function. The corolla is localized in the periphery of the IS where active TCR-CD3 is initiated^35, 40, 41^. Incubation of T-cells on SLB with only CD58 results in CD2 microdomains that are enriched in signaling molecules associated with TCR signaling whether or not the TCR is directly engaged^7, 42^. It is reported that the cytoplasmic tail of CD2 is required for this enrichment of signaling molecules and we report here that it is also needed for efficient corolla formation. We, thus envisioned, that the CD2 corolla is harnessing this peripheral signaling domain to amplify TCR signals hence boosting T-cell activation.

We investigated whether the corolla recruits signaling molecules and how this compares to TCR microclusters. Staining for active phosphorylated Src kinases (pSrc), phosphorylated PCLγ1 (pPLCγ1) and phosphorylated LAT (pLAT) at 15 min of IS formation displayed significant enrichment in the corolla (Figure 5a-c). In the absence of CD2 ligation, pSrc, pPLCγ1 and pLAT clusters were primarily found associated with peripheral TCR microclusters, which have a much smaller area and lower intensity, compared to signals in the corolla. Quantification of the integrated pSrc, pPLCγ1 and pLAT signals revealed 2-3-fold increase in total signal due to the corolla compared to TCR microclusters formed with ICAM1 and anti-CD3 Fab’ only (Figure 5d). This was also confirmed using antigen-specific bilayers with human T-cells (data not shown). Thus, CD2-CD58 interactions in the corolla can amplify early TCR signaling.

**Figure 5.**
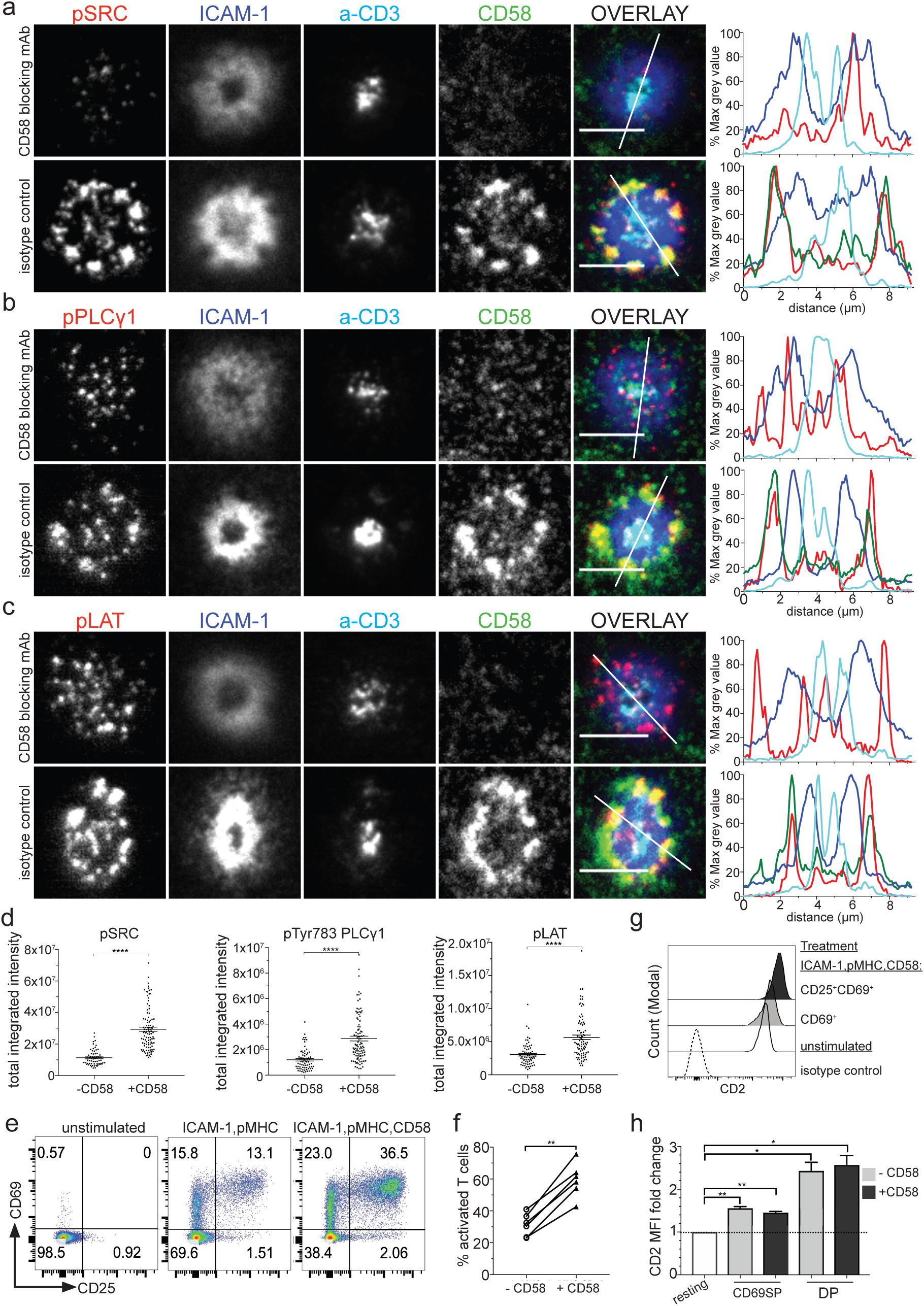
CD2 corolla amplifies TCR and LAT dependent signals and enhances T-cell activation. Representative images of CD8^+^ T-cells incubated on ICAM1 (200/μm^2^), anti-CD3 Fab (30/μm^2^) and CD58 (200/μm^2^) reconstituted SLBs, fixed 15 min post-incubation, followed by permeabilization, blocking and intracellular staining for phosphorylated Src kinases **(a)**, pTYR783PLCγ1, pPLCγ1 **(b)** or pTYR171LAT, pLAT in **(c).** SLBs were previously incubated with blocking anti-CD58 mAb (TS2/9 clone) or an isotype control. Same results were obtained when CD58 was omitted instead of using CD58 blocking mAb. Cells were imaged with TIRFM. Scale bar, 5 μm. Histograms depict the intensity profiles on the diagonal white lines in overlay images. y-axis; raw pixel intensity signal normalized to maximum intensity pixel of each channel. The CD58 signal was omitted in CD58 blocking conditions for clarity. **d)** Levels of pSrc, pPLCγ1 and pLAT for an average of 80 cells per condition shown in (a-c). ****, p value <0.0001 with non-parametric Mann Whitney *U-*test. A representative of three experiments is shown. Error bars, SEM. **e)** Representative plots, from 1 of 3 donors, showing the activation status of *1G4*^+^ TCR memory CD8^+^ T-cells after 24 hrs culture in resting conditions or on SLBs presenting ligands of two different compositions. **f)** The percentage of activated (CD69 single positive, and CD69^+^CD25^+^) 1G4^+^ T-cells after treatment of cells as in (e), for 6 donors tested. **, p value <0.007 with non-parametric Mann Whitney *U-*test. **g)** Representative histograms of CD2 expression, from 1 donor, in resting (solid empty), activated CD69^+^ single positive (grey), activated CD69^+^CD25^+^ (black) T-cells; isotype-control stained (dashed empty histogram). **h)** From experiments in (e), the mean fold change in CD2 mean fluorescence intensity, MFI, from 3 donors, for CD69^+^, single positive (CD69SP) and CD69^+^CD25^+^, double positive (DP) activated T-cells in the presence (+) and absence (-) of CD58, compared to resting conditions. **, p>006; *, p>0.019 with *U*-test and Welch’s correction.

We next asked if the CD2 corolla-mediated enhancement of proximal TCR signaling results in augmented T-cell activation. We, thus, tested T-cell activation 24 h post-incubation on ligand reconstituted SLBs with and without CD58. We used *1G4* TCR transfected human memory CD8^+^ T-cells. We detected an increased percentage of activated T-cells and expression of T-cell activation markers (CD69 and CD25) in response to SLB presenting pMHC in the context of ICAM1 when CD58 was also present (Figure 5e-f). CD2 expression was also increased in activated T-cells in both the presence and absence of CD2 ligation (Figure 5g-h); CD69^+^CD25^+^ T-cells showed a 2.5-fold increase in CD2 mean fluorescence intensity (MFI) compared to unstimulated cells followed by CD69^+^CD25^−^ T-cells with a 1.5-fold increase in CD2 MFI. This implies a positive feedback effect of CD2 on enhanced TCR signaling over hours of stimulation, by means of enhancement of its own surface expression to increase signaling domain area. Our results suggest that CD2 corolla formation is yet another intrinsic property of CD2 ligation that contributes to its costimulatory function via a spatial effect that favors enhancement of TCR signaling.

### CD2 expression levels scale TCR signaling

We have shown that CD2 levels and the cytoplasmic tail of CD2 are key determinants in CD2 corolla formation and that CD2-CD58 interactions boosts TCR signaling in the IS. We, therefore, sought to determine how CD2 levels influence quantitative aspects of CD58 engagement and CD2 signaling. hCD2 with a C-terminal mRuby was transfected into AND T-cells and left to interact with ICAM1, pMHC and CD58 reconstituted SLBs for 15 min. There was a strong positive correlation between intensity of the hCD2 and CD58 signals (Figure 6a). This supports a model of 1:1 interaction of laterally mobile and monodispersed CD2 and CD58 driving their mutual accumulation in growing microdomains. However, this measurement does not take into account the total cellular CD2 due to the TIRF excitation. We, further, tested the relation between total hCD2 expression and CD58 accumulation by testing separate populations of AND T-cells expressing increasing levels of total cellular hCD2. This confirmed a stronger accumulation of CD58 in the IS with higher cellular CD2 expression levels (Figure 6b). More importantly, increasing levels of CD58 accumulation corresponding to increasing levels of CD2 expression, correlated with integrated pSrc (Figure 6c). Intriguingly, the intercept of the regression line, for this plot, with the Y-axis corresponds to the level of pSrc observed in the absence of CD58 (Figure 5d and 6c). Therefore, the total number of hCD2 molecules expressed on each cell controlled the increase in pSrc in a linear fashion. It is consequently, fair to propose that exhausted TILs exhibiting the CD2^low^ phenotype will suffer from a lower CD2-mediated enhancement of TCR signalling upon tumour antigen recognition.

**Figure 6.**
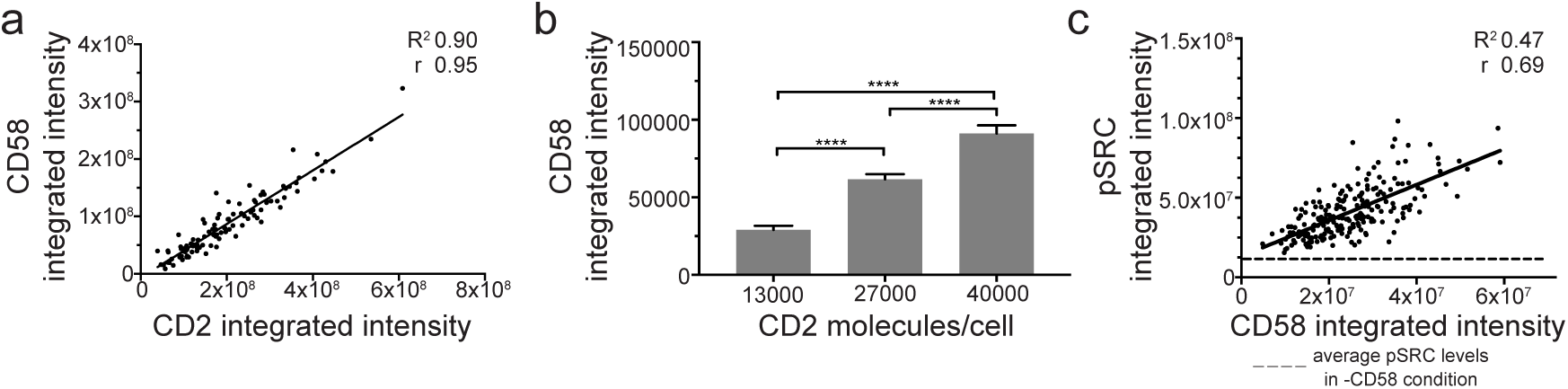
CD2 expression levels scale TCR signaling. **a)** AND T-cells were transfected with hCD2FL-Ruby at levels similar to memory CD4^+^ T-cells. Cells were plated on ICAM1 (200/μm^2^), pMHC (30/μm^2^) and CD58 (200/μm^2^) containing SLBs for 15 min before fixation and imaged with TIRFM. Shown, here, is the intensity of CD2FL (x-axis) and CD58 (y-axis) accumulating in each single T-cell immune synapse (~100 cells from 1 experiment). ****, p<0.0001; there is a positive correlation of Pearson r ~0.95, R^2^ 0.90. A representative of three experiments is shown. **b)** AND T-cells were transfected with increasing levels of hCD2FL-Ruby and treated as in (a). The graph shows the average intensity of CD58 accumulation for each different AND T-cell population (~100 cell/population). Error bars, SEM. **c)** CD8^+^ T-cells were incubated on ICAM1 (200/μm^2^), anti-CD3 Fab (30/μm^2^) and CD58 (200/μm^2^) reconstituted SLBs for 15 min before fixation, followed by permeabilization, blocking and intracellular staining for phosphorylated Src kinases, pSRC. There is positive correlation between the two parameters with Pearson r 0.69; R^2^ 0.47; ****, p<0.0001; y-intercept 1.32×10^7^, slope 1.123;- - - -, dashed line represents the mean pSRC levels (1.15×10^7^) from a population of single T-cells, activated as above but in the absence of CD58.

### High PD-1 levels inhibit CD2-mediated enhancement of TCR signaling during synapse formation

We showed that CD2 ligation results in localization of CD2 in the periphery that allows enhancement of TCR proximal signaling and T-cell activation in response to antigen (Figure 5–6). We have correlated corolla formation and CD2-mediated enhancement of TCR signals with CD2 expression levels. We thus suggest that CD127^low^PD-1^hi^ TILs from cancer patients could have a compromised CD2 pathway, due to their CD2^low^ phenotype. This could affect their overall activation potential due to weaker TCR enhancement and in combination with their high expression of inhibitory checkpoint receptors, may contribute to worse patient prognosis.

We, finally, asked whether the high levels of PD-1 expressed in exhausted TILs could impact CD2 signaling. We investigated the impact of high PD-1 on CD2-mediated enhancement of pSrc in the IS of memory CD2^hi^ CD8^+^ T-cells from peripheral blood of healthy individuals. Our absolute quantification of PD-1 levels in exhausted TILs allowed us to reproduce these levels, using mRNA electroporation, in these CD2^hi^ CD8^+^ T CD8^+^ T-cells. Electroporated PD-1^hi^CD2^hi^ memory CD8^+^ T-cells were incubated on SLBs reconstituted with ligands of four different compositions (Figure 7). CD2-CD58 and PD-1-PD-L1 interactions were colocalized in the CD2 corolla formed by CD8^+^ memory T-cells expressing PD-1 (Figure 7a). CD2 ligation significantly boosted pSrc signals and additional ligation of PD-1 resulted in a significant 20% reduction of CD2-mediated enhancement of TCR signalling (Figure 7b). There was no detectable PD-1 effect on pSrc signals in the absence of CD2 ligation. This suggests that PD-1 targets CD2-mediated enhancement of TCR signaling rather than TCR proximal signalling in this setting. Moreover, in half of the patients, >50% of CD127^−^PD-1^+^ CD8^+^ TILs were CD28 negative too (Supplementary Figure 5). CD4^+^ TILs, did not suffer the same loss of CD28 as CD8^+^ TILs. In the absence of CD28, the CD2^low^ phenotype either by itself or in addition to PD-1-mediated inhibition of CD2-enhancing property on TCR signaling could become an even stronger limiting step for T-cell activation and effective T-cell functions. In summary, we showed that high PD-1 levels, a feature of exhausted TILs, can significantly reduce CD2-mediated enhancement of TCR signaling when CD2 is expressed at normal levels. Therefore, exhausted TILs exhibiting the CD2^low^ phenotype would be even more susceptible to the negative outcome of PD-1-mediated inhibition of CD2-induced effects.

**Figure 7.**
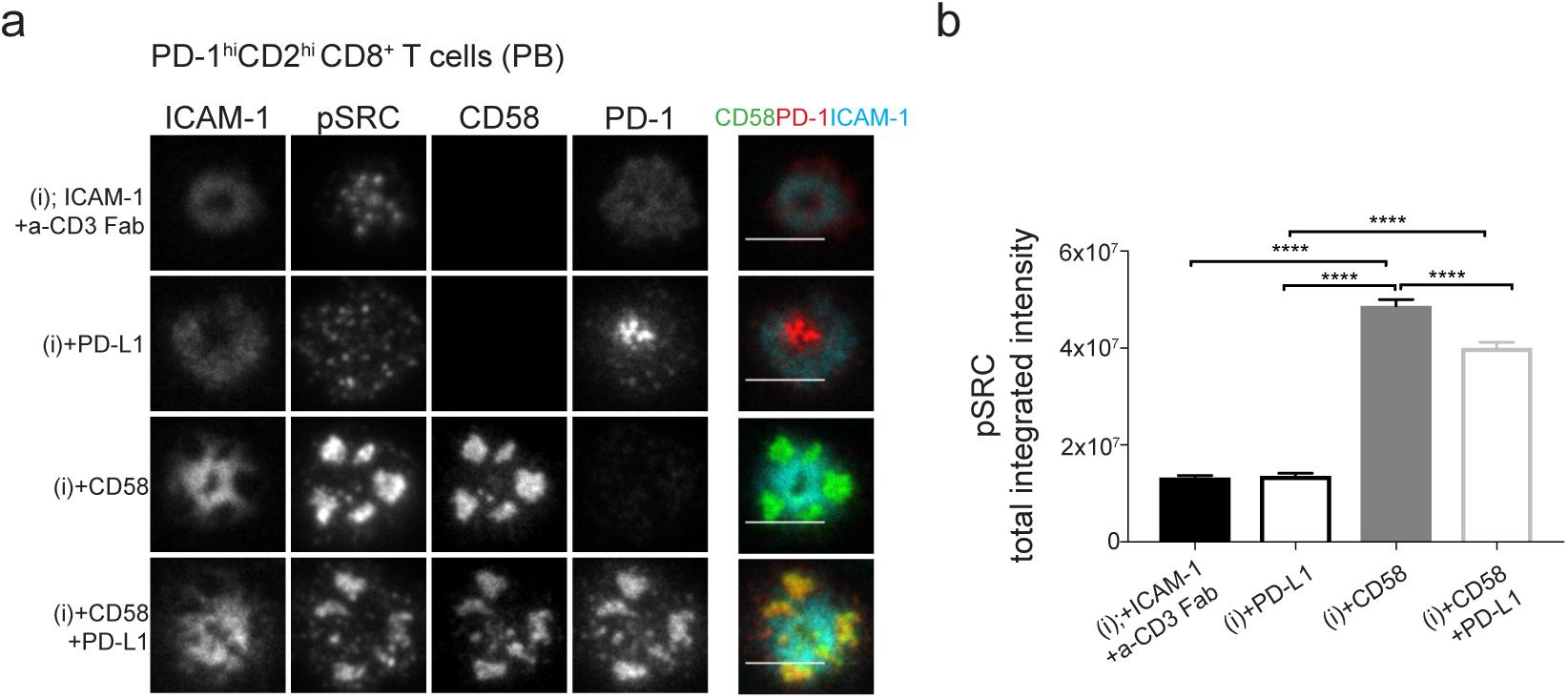
High PD-1 levels inhibit CD2-mediated enhancement of TCR signaling during synapse formation. **a)** Representative images of memory CD2^hi^CD8^+^ T-cells, isolated from peripheral blood of healthy individuals, incubated on four different ligand compositions based on ICAM1, anti-CD3 Fab (i), ±CD58, ± PD-L1 reconstituted SLBs, fixed 15 min post-incubation, followed by permeabilization, blocking and intracellular staining for phosphorylated Src kinases (pSRC) and imaged with TIRFM. Cells were transfected to express high levels of fluorescently-tagged surface PD-1 (~1.5×10^4^ molecules/cell) i.e. PD-1^hi^ similar to CRC CD8^+^ TILs. Scale bar, 5 μm. **(b)** pSRC levels at the IS of single memory PD-1^hi^ CD2^hi^CD8^+^ T-cells from experiment in (a). At least 100 cells per condition where analysed. Representative data from one out of three donors tested is shown. ****, p>0.0001 with non-parametric Mann Whitney *U-* test. Error bars, SEM

## Discussion

Using model systems for T cell activation (T:APC cell conjugates and T:SLB conjugates) we revealed unexpected properties of CD2 that contribute to its function as an adhesion receptor and positive costimulatory molecule. We show that engagement of sufficient CD2 numbers with its natural ligand results in a unique protein rearrangement in the IS, the peripheral “corolla” pattern in the dSMAC, that enhances TCR signaling and uniquely changes the localization of classical co-stimulatory/checkpoint receptors, CD28, ICOS and PD-1. Kaizuka *et al* previously reported segregation of CD2 and TCR upon engagement of both receptors by laterally mobile ligands on SLBs^7^, but the use of T:APC conjugates and the presence of ICAM1 in our SLB system allowed us to fully appreciate the protein organisation of CD2-CD58 microdomains in the IS and further investigate their relationship with other receptors at the IS. We propose that CD2 corolla effectively enhances TCR signaling by sustaining CD2 clusters in the periphery of the IS where signaling is initiated^35, 41, 43^, in contrast to terminating signaling by translocation to extracellular vesicle in the synaptic cleft^44^. TCR microclusters traverse the corolla, but TCR proximal signaling seems to spread through the entire corolla area. Similarly, capturing of costimulatory/inhibitory molecules by the corolla sustains the receptors in the periphery potentially enhancing their signaling output. Peripheral engagement of CD28 was shown to boost CD28-mediated enhancement of IL-2 production compared to a centrally-localized CD28 signal colocalizing with anti-CD3 stimulation^45^. Moreover, CD2 corolla could function as an expanded adhesion area of the IS across which a T-cell can extend its scanning activity of the APC surface.

Corolla formation displays a number of unexpected features – notably the persistence of thousands of CD2-CD58 interactions in the dSMAC, in the face of centripetal F-actin flow. CD2-CD58 interactions are transient with a half-life of only a fraction of a second^46^. Due to the high lateral mobility of CD2, most CD2 molecules on the cell can diffuse into the IS and engage CD58. Since CD2-CD58 interactions maintain a density of ~800 μm^2^ as the number of interactions increases, cells with 4×10^4^ CD2 molecules will have total CD2 microdomain area approaching 50 μm^2^, which is about half the area of a mature IS^47^. Thus, when TCR-pMHC and LFA-1-ICAM1 interactions are transported inward by strong links to F-actin, the freely mobile CD2 protein and its interaction partner CD58 may be forced into the distal aspects of the IS by competition for space as the microdomains grow to an extent where the only available area is in the distal compartment. This turns out to be a strategic position for CD2 as it is the origin of TCR microclusters that fuel IS and corolla formation. The cytoplasmic domain of CD2 promoted corolla formation, but it’s not clear if this is through a change in the adhesion area, or a feed forward effect related to the TCR-mediated increase of CD2 avidity for its ligand^48, 49, 50^, which is lost without the cytoplasmic tail. The cytoplasmic tail also links CD2 via CD2AP and CIN85 adaptor proteins to other signaling proteins and to the actin cytoskeleton^51^, another potential key player in driving efficient corolla formation.

In light of the CD2-level-mediated scaling of TCR signalling we propose that the CD2^low^ phenotype renders TILs, and in particular exhausted CD8^+^ TILs, less responsive to tumour antigens and may compromise tumour clearance. It has been shown that PD-1-mediated inhibition preferentially targets CD28-mediated costimulation^19^. We have identified here a new pathway for PD1-mediated inhibition in T-cells, whereby even in the absence of CD28 engagement, PD1 can also attenuate CD2-mediated costimulation, hence limiting T-cell activation and function. The amount of PD-1 expression regulates the magnitude of PD-1-mediated inhibition of T-cell activation and cytokine production^52, 53^ hence PD-1 acts a quantitative checkpoint. Similarly, we have shown that CD2 levels regulate corolla formation and CD2-mediated enhancement of TCR signalling hence CD2 expression can act as a quantitative checkpoint for synapse organisation and T-cell activation. From our findings, we can predict a 30% reduction in CD2-mediated boost of TCR in CD2^low^ exhausted CD8^+^ TILs. This CD2 molecular deficit could act as a limiting parameter for this checkpoint that will be further compromised by a 20% mediated by the PD-1 pathway.

In addition, CD28^−^CD8^+^ T-cells are enriched in the periphery of older adults and are CD2^hi^ (CD2 memory levels) similar to CD2^hi^CD28^+^ CD8^+^ T-cells^54^; it was reported that CD28^−^CD8^+^ T-cells use CD2 as their primary costimulatory pathway during T-cell activation^55^. We show that CD8^+^ TILs are often CD28^−^ and instead of showing high CD2 levels, they exhibit a CD2^low^ phenotype. This highlights further the potential impact that the CD2^low^ phenotype can have on exhausted TIL function.

Some of our results suggest that this CD2^low^ phenotype is also detectable at the mRNA level, but this does not exclude other post-translational regulation. Why TILs from some CRC patients retain high levels of CD2 expression that are commensurate with their being activated effector T-cells is yet to be addressed. Monitoring CD2 level changes upon treatment of CRC patients with anti-PD-1 mAbs that can result in reinvigoration of circulating exhausted CD8^+^ T-cells is an important line of future study. We have shown that successful T-cell activation results in upregulation of CD2 expression. Further investigation will reveal if the CD2^low^ phenotype is specific to CRC or present in other types of solid tumours. Finally, the CD2^low^ phenotype may also be used as an indicator of poor prognosis or a predictor of poor response of CRC patients to certain immunotherapies and could be, by itself, an interesting target for immunotherapy. Harnessing these two checkpoints i.e. by blocking high PD-1 levels and restoring CD2 deficit could more efficiently unleash an effective and more sustained anti-tumour response by exhausted TILs.

## Acknowledgements

We thank S. Davis, P.A. Van Der Merwe, O. Dushek for kindly providing plasmids and HLA-A2 pMHC monomers. M. H. Brown, for helpful discussion and feedback on experiments and writing of manuscript and all the Dustin lab members for their kind support. We thank S. Balint and D. Depoil for maintaining the TIRF microscope and C. Laggerholm for access to and assistance with the Airyscan confocal microscope. We thank S. Sivakumar for his helpful discussion, support and feedback on patient data. A Kennedy Trust for Rheumatology (KTRR) Prize Studentship supported P.D. An UCB-Oxford Post-doctoral Fellowship supported EAS. Wellcome Trust Principal Research Fellowship 100262Z/12/Z and a grant from KTRR supported M.L.D. and E.A.S. KTRR supports the Kennedy Institute Microscopy Facility and the Wolfson Foundation supports the Weatherall Institute Microscopy Facility. A collaborative grant from the Human Frontiers Science Program supported E.A.S., A.S. and M.M.H. A fellowship from Philippe Foundation partially supported D.D.. A Wellcome Trust Senior Research Fellowship 207537/Z/17/Z supported O. D., E.A.S., and M.K.. We acknowledge the contributions the Oxford Gastro-Intestinal Biobank and the Oxford Inflammatory Bowel Disease Cohort study, which are supported by NIHR Biomedical Research Centre, Oxford. The views expressed are those of the authors and not necessarily those of the NHS, the NIHR or the Department of Health.

## Author Contributions

P.D. conceptualised the project, designed and performed experiments, analysed the data and co-wrote the manuscript. E.A.S. designed, performed and analysed experiments. S.V. prepared and performed experiments and maintained critical infrastructure. K.K. assisted in and acquired confocal microscopy experiments. S.M., M.F., E.M., E.A.S., prepared single cell suspensions from CRC tissue. S.M. performed transcriptional analysis. P.D., E.A.S, performed the staining and acquisition experiments of CRC tissues. J.A., S.V., V.M., M.K. prepared essential reagents for experiments. V.M. designed image analysis software and trained P.D. in its use. L.L. and T.S. performed transcriptional analysis of CRC cohorts. The O.IBDC.I. provided access to CRC tissue and clinical data. P.D., E.A.S., V.M., D.D., A.S., M.M.H. and M.L.D. made intellectual contributions to the project through regular discussions. D.D. provided training to P.D., conceptualised and designed experiments. M.L.D. supervised the research and facilitated collaboration. P.D. drafted the manuscript and E.A.S., V.M., D.D., and M.L.D. contributed to writing and editing.

## Methods

### Reagents

RPMI 1640 (31870074), Dynabeads Human T-Activator CD3/CD28 for T-cell expansion and Activation kit (1132D) were purchased from Thermofisher Scientific. Hyclone Fetal bovine serum was obtained from Fisher Scientific. Poly-L-lysine solution 0.1% (w/v) in H_2_O was obtained from Sigma (25988-63-0). The following anti-human antibodies were purchased from Biolegend, CD62L (DREG-56), CD3 (UCHT-1), CD4 (A161A1), CD8 (HIT8a), CD2 (RPA2.10), CD28 (37.51), CD28 (CD28.2), PD-1 (29F.1A12), TCRα/β (IP26), CD11a (TS2/4), mouse IgG1 isotype control (MOPC-21), CD58 (TS2/9), mouse IgG1 isotype control (MG1-45), CD45 (HI30), CD127 (A019D5), HLA-A2 (BB7.2), CD45RO (UCHL1). The following anti-human antibodies were purchased from BD Bioscience, CD45RA (HI100), CD127 (HIL-7R-M21), CD58 (1C3), CD2 (CD2.1) clone was a gift of D. Olive (Aix Mareseille University). The following antibodies were purchased from Cell Signaling, pPLCγ1(pY783), pLAT(pY171), pSRC (pY416) (D49G4). The following anti-mouse antibodies were purchased from Biolegend, CD4 (RM4-5), CD3 (145-2C11), TCRβ (Η57-97). TCRβ (Η57-97) Fab was obtained from Bio X Cell.

### TCGA human cohort data

The human cohort data was obtained from The Cancer Genome Atlas (TCGA) which is publically available for the scientific research community, and comprises the transcriptomic profiles of 255 CRCs in the form of RSEM count data at the mean. Ethical approval for the data was obtained from the National Institute of Health (NIH). Expression data from the TCGA colorectal cancer cohort was obtained directly from the Broad Institute website (https://gdac.broadinstitute.org).

### Metastatic lymph nodes in colorectal cancer (METALLIC) cohort

Metastatic lymph nodes in colorectal cancer (METALLIC) cohort biospecimens were collected from newly diagnosed patients with stage III sporadic colon adenocarcinoma who had undergone surgical resection and had received no prior treatment for their disease at the Oxford University Hospitals between March 2012 to May 2015 (n=148). All patients had given consent for this study and this was performed under ethics reference 11/SC/0236. Cases were staged according to the *European Society of Medical Oncology (ESMO)* guidelines. Patients were excluded from this analysis if there was a previous history of CRC, any predisposing conditions for CRC (such as IBD, polyposis syndromes), patients presenting with synchronous CRC lesions or if there was evidence of inherited CRC predisposition syndromes.

### Data analysis – CD8 exhausted T cell subpopulations

Gene expression of different CD8 exhausted T cell subpopulations in the count data from the TCGA and METALLIC cohorts was analysed using publically available genesets from the molecular signatures database (MSigDB) maintained by the Broad Institute (GSE 41867 genesets). Gene expression of different cell subpopulations was assessed from transcriptomic data by calculating a mean value for expression of each geneset in each sample.

Statistical analyses were performed for each CD8 exhausted T cell geneset in R version 3.5.2. Linear regression models, generated using the *glm* function in R, were utilised for testing for statistical significance between CD8 exhausted T cell geneset expression and CD2 gene expression. Marginal plots were generated using the ‘ggplot2’ and ‘ggExtra’ r packages (*ggMarginal*).

### Isolation of T-cell subsets, cell culture and T-cell blast generation

Human T-cell subsets were isolated from leukapheresis produts using negative selection kits from STEMCELL Technologies. Rosette Sep (Stemcell Technologies) was used to enrich for total CD8^+^ or CD4^+^ T-cells. Naïve and memory cells were isolated using EasySep Enrichment kits (negative selection, Stemcell Technologies). Flow cytometric analysis based on CD62L and CD45RAwas used to confirm purity is above >92%. Human T-cells were cultured in resting conditions, with no IL-2, at 37°C, 5% CO_2_, in RPMI 1640 (Roswell Park Memorial Institute) medium (Life Technologies) supplemented with 10% FBS, 5% Penicillin-Streptomycin, Penstrep (Gibco), 1× MEM Non-Essential Amino Acids Solution (Thermo Fisher Scientific), 10mM HEPES (Life Technologies), 1mM Sodium Pyruvate (Thermo Fisher Scientific). T-cells were cultured at 2×10^6^ cells/ml for a maximum of 3 days. Human T-cells isolated were also used to generate T-cells blasts using the Dynabeads Human T-Activator CD3/CD28 for T-cell expansion and Activation kit (Life Technologies) as per company’s protocol. Human T-cells blasts were ready to be used in experiments on Days 6-7 of expansion protocol. No IL-2 was supplemented in media the day before an experiment.

AND Mouse T-cells (AND blasts) were generated from spleen and lymph nodes of 2-3 months old AND B10.Br mice; mice were housed under pathogen-free conditions in the Kennedy Institute of Rheumatology Animal Facility in accordance with local and Home Office regulations. Briefly, spleen and lymph nodes were crashed over 70 μm strainer to generate single cell suspensions and following red blood cell lysis, incubated with 1 μM moth cytochrome c, MCC, peptide and 50 U recombinant human IL-2 in High Glucose (25mM) Dulbecco’s Modified Eagle Medium supplemented with 0.116g/L L-Arg-HCl, 0.036g/L L-asparagine, 2g/L NaHCO_3_, 2.385 g/L HEPES, 1mM sodium pyruvate, 1.5 mM L-glutamine, Pen/Strep and 50 μM 2-mercaptoethanol at 37°C, 5% CO_2_ incubator. AND T-cells were used for experiments on day 6-7 of culture; cell culture was enriched (>95%) in AND CD4^+^ T-cells. No IL-2 was supplemented in media the day before an experiment.

### Cell line culture

The CF996 EBV-transformed B cell line (Sigma Aldrich), was maintained in supplemented RPMI-1640 medium with 10 % FBS at cell densities of 0.75-1×10^6^ cells/ml in a 37°C, 5% CO_2_ incubator.

### Generation of monocyte-derived DCs

Human monocytes were isolated from leukapheresis products using the Rosette Human Monocyte Enrichment Cocktail kit (Stem Cell). Purified cells were cultured in 2 ml/well in 12 well plates for at least 4 hr at 37°C to allow for adhesion and enrichment of monocytes. Culture supernantant was resuspended and removed. Cells were washed twice with fresh media before adding culture media supplemented with 0.1 μg/ml, granulocyte-monocyte colony stimulating factor, GM-CSF (Immunotools), and 0.05 μg/ml IL-4 (Peprotech). Cells were incubated for 24 hr in the above conditions at a 37°C and 5% CO_2_ incubator. Culture supernatant was supplemented the day after with a final concentration of 0.1 μg/ml prostaglandin E, 0.04 μg/ml TNF-α, 0.02 μg/ml IL-1β and 0.02 μg/ml IFNγ. This 24hr incubation was used to activate cells.

### RNA transfection of human and mouse T-cells

RNA constructs were prepared from DNA constructs in pGEM plasmids linearised with SpeI restriction enzyme at 37°C for 1-2 hrs. DNA precipitation, *in vitro* RNA transcription and *in vitro* poly-adenylation were performed exactly as described in the protocol of the mMESSAGE mMACHINE T7 ULTRA Transcription kit (Thermofisher Scientific). RNA was purified as per instructions of MEGAclear Transcription Clean-up kit (Thermofisher Scientific). RNA was finally aliquoted at 5 μg per PCR/vial and stored at −80°C freezer. RNA was only thawed once and used immediately.

Freshly isolated T-cells (human or mouse) or T-cells in culture (human or mouse) were harvested and washed three times in Opti-MEM media (LifeTechnologies). Depending on the expression levels to be achieved and the number of available cells, T-cells were resuspended between 1×10^6^-2.5×10^6^ per 100 μl. Each 100 μl cell suspension was mixed with the RNA preparation and transferred to a Gene Pulser/Micropulser electroporation cuvette, 0.2 cm gap (BIO-RAD). Cells were electroporated at 300V, 2ms in an ECM 830 Square Wave Electroporation System (BTX). Cells were harvested using 0.5 ml of pre-warmed culture media at 37°C and cultured at ~1-2×10^6^ cells/ml in warm media. Cells were incubated for at least 12 hr in 37°C, 5% CO_2_ incubator before testing expression levels and transfection efficiency. For TCR RNA transfection, as described in^29^, we used 5 μg of each TCRα-β chain and 5 μg of CD3-ζ chain for 2-5×10^6^. BTX cuvettes, 2mm gap (VWR) were used for TCR transfection. Two TCR constructs were used: the *6F9* TCR that recognizes an HLA-DPB1*04:01-restricted MAGE-A3 melanoma antigen^56^ and the *1G4* TCR that recognizes an HLA-A*0201-restricted NY-ESO-1 cancer-testis antigen^57^.

### Supported Lipid Bilayer (SLB), immunostaining and TIRFM Imaging

SLB were prepared as described previously in^44, 58^. Briefly, glass coverslips (Nexterion) were cleaned in piranha solution, rinsed in water, dried, plasma cleaned and mounted onto six-channel chambers (Ibidi). Small unilamellar liposomes were prepared using an extruder (Avestin, 100 nm pore size filter) using 18:1 DGS-NTA(Ni), NTA-lipids, (Stratech, Avanti Polar Lipids, 790404C-AVL) that bind HIS-tag proteins, CapBio 18:1 Biotinyl Cap PE (Stratech, Avanti Polar Lipids, 870282C-AVL), CapBio-lipids, and 18:1 (Δ9-Cis) PC, DOPC-lipids, (Straterch, Avanti Polar Lipids, 850375C-AVL). Lipids were prepared in stock concentrations of 25% (4mM) NTA-lipids, 4mM CapBio-lipids and 4mM DOPC-lipids. For experiments NTA-lipids were used at a final concentration of 12.5% and CapBio-lipids were used at a concentration determined to give 30 molecules/μm^2^ of monobiotinylated anti-CD3 Fab or monobiotinylated peptide MHC. Channels in Ibidi chamber cover with liposome mixture and after a 20 min incubation at RT, they were washed with human serum albumin (HSA)-supplemented HEPES buffered saline (HBS) supplemented with 1 mM CaCl_2_ and 2 mM MgCl_2_, referred to as HBS/HSA. SLBs were blocked with 5% casein in PBS containing 100 μM NiSO4, to saturate NTA sites and washed with HBS/HSA. Unlabelled/labelled streptavidin was added to channels coupling of biotinylated proteins to cap-biotinyl-containing lipids. Combination of the following reagents: Biotinylated UCHT1 Fab (30 molecules/μm^2^), peptide-loaded HLA-A2, HLA-DP4 (30 molecules/μm^2^), His-tagged ICAM1 (200 molecules/μm^2^), CD58 (200 molecules/μm^2^), CD80 (molecules/μm^2^) ICOSL (100 molecules/μm^2^), PD-L1(100 molecules/μm^2^), moth cytochrome c peptide (MCC), MCC-loaded I-E^K^ (0.1 or 30 molecules/μm^2^), were then incubated onto the SLBs at specific concentrations to achieve the indicated molecule densities. Channels were washed and kept in HBS/HSA until ready to be used. 6F9 TCR transfected human CD4^+^ T-cells or AND T-cells were stained with anti-TCRβ (Η57) Fab to allow tracking of TCR during imaging. For fixed experiments, cells were allowed to interact with SLBs for 15 min at 37°C, before a 10 min fixation with 2% PFA in PHEM buffer at 37°C. Cells were permeabilised with 0.1% Triton X-100 in HBS/HAS for 2 min at RT where required. Cells were blocked in 5% Casein containing 5% Donkey serum for 60 min at RT. Cells were stained overnight at 4°C with primary anti-pSRC, anti-PLCγ1, anti-pLAT prepared in 5% cazein containing 5% donkey serum followed by washing and incubation in secondary donkey anti-rabbit antibody for 45 min at RT. Cells were immediately imaged or fixed as above for 5 min at RT and stored at 4°C. Imaging was performed on an Olympus IX83 inverted microscope equipped with a TIRF module. The instrument was equipped with an Olympus UApON 150× TIRF N.A 1.45 objective, 405 nm, 488 nm, 568 nm and 640 nm laser lines and Photomertrics Evolve delta EMCCD camera. In live imaging experiments, SLBs were transferred to a pre-heated incubator on top of the TIRF microscope and cells were added in the well.

### Cell conjugates and imaging

CF996 EBV-transformed B cells were stained with 0.5 μM cell trace violet (CTV, Thermofisher) by incubation at 37°C, 5% CO_2_ for 10-15 min. 10% FBS was added to quench dye and cells washed twice in media. MAGE-A3_243-258_ or NY-ESO-9V_157-165_ loaded CF996 EBV-transformed B-cells were conjugated with TCR transfected CD4^+^ and CD8^+^ T-cells at 1:1 ratio, in round bottom tubes for 25-30 min at 37°C. Conjugates were transferred onto a poly-L-lysine coated μ-slide 8-well glass bottom chambers and fixed with 4% paraformaldehyde for 10 min at 37°C. Washing steps with PBS/0.5%BSA were performed in-between each of the following steps. Cells were washed, permeabilised for 4-5 min with 0.1% Triton-X100 prepared in PBS/0.5%BSA for 10 min and blocked with PBS/5%BSA/5% donkey serum for 1hr at RT. Cells were washed and incubated with unlabeled, non-blocking anti-CD2 (CD2.1) in PBS/5%BSA, overnight at 4°C. Cells were washed and incubated in secondary antibody, AF647-conjugated donkey anti-mouse (Jackson ImmunoResearch Laboratories) for 45 min at RT. Cells were washed and fixed with 2% PFA for 5 min at RT. Cells were washed and incubated in FITC-conjugated anti-CD11a (TS2/4) in PBS/0.5%BSA for 1hr, at RT. Cells were washed and fixed wit 2% PFA for 5 min at RT. Finally, cells were loaded with Vectashield antifade mounting medium (H-1000) to minimize the movement of cell conjugates during imaging. Imaging was performed on a Zeiss inverted LSM 880 AiryScan confocal microscope in xyz dimensions in the airyscan mode. We acquired the Airyscan data using the ZEISS ZEN software used to operate the microscope and the airyscan settings were used in the automated mode. The instrument was equipped with a Plan-Apochromat, 63×, NA. 1.40 oil objective.

### Activation assay on SLBs

Ligand reconstituted SLBs were prepared as described above. ~ 0.5×10^6^ human memory CD8^+^ T-cells, transfected to expressed *1G4* TCR as described above, were incubated on SLBs per channel, in supplemented RPMI and incubated for 24hr at 37°C, 5% CO_2_ incubator. Cells were, then, harvested using ice-cold PBS/0.5%BSA, washed and stained for flow cytometry analysis of their activation status as described below.

### Flow cytometry analysis to determine ligand and receptor expression levels

1×10^6^ peripheral blood mononuclear cells, PBMCs, or 10^5^ cells (T-cells, CF996 B-cell line, monocyte or DCs) were stained in 96-well U-bottom plates. For determination of activation status of memory CD8^+^ T-cells, activated on SLBs, ~0.25×10^6^ cells were stained. Cells were stained in fixable viability dye efluor 780 (eBioscience) in ice-cold PBS/0.5%BSA at 4°C for 10 min. A washing step with PBS/0.5%BSA was carried in-between each of the following steps. For staining PBMCs, monocytes, DC and B cells, an Fc block step was performed by resuspending cells in 50μl of Fc block prepared in PBS/5%BSA (room temperature) using 5μl of Human Trustain FcX (Biolegend) per sample i.e. 5μl/50μl of PBS/0.5%BSA. Cell were incubated at room temperature for 10 min. Cells were stained with an antibody mix prepared in ice cold PBS/0.5%BSA (and Fc block in the case of PBMCs, monocytes, DC and B cells) and incubated on ice for 15-20 min. Cells were then ready to be analyzed. Cells were analysed on an LSRII or Fortessa machine. Analysis of acquired data was performed on FlowJo 10.4.2 and GraphPad Prism 7. For absolute quantification of CD58, CD2 and PD-1 expression, the Quantum Simply Cellular anti-mouse IgG kit was used (Bangs Laboratories, see further details in next section). We counted two CD2 molecules for each anti-CD2 IgG detected as we have previously found that the anti-CD2 mAbs bind bivalently at 10 μg/ml^59^. We applied the same for CD58 and PD-1 quantification. For CD58 densities on the B-cell line, we determined a mean surface area 277.6 μm^2^ using the CASY cell counter and analyser.

### Determining absolute molecular densities on SLB-coated silica beads and cells

Silica microspheres (SS05003/SS05N) (Bangs Laboratories) were used for SLB preparation. In order to simulate the surface area used in our SLBs prepared in 6-well chamber IBIDI slides, we used the number of beads that gives us the same surface area as the surface area covered by the SLBs in Ibidi channels. Silica bead vial was vortexed briefly to uniformly resuspend silica beads. The required volume of beads was transferred to a clean 1.5ml Eppendorf tube and 1 ml of PBS was added to wash the beads by spinning beads down. This was repeated twice. Beads were then incubated in the required mixture of lipids as described in ‘Supported Lipid Bilayer (SLB), immunostaining and TIRFM Imaging’ and the same steips were performed from this point onwards. If titration of biotinylated proteins was used, cap-bio lipids were titrated instead of the protein itself, hence silica bead with different lipid mixtures were prepared and processed separately. If biotinylated proteins were titrated, streptavidin, at saturating concentrations, was added to the silica beads for 10 min at RT. His-tag proteins were serially diluted and added in 100 μl of a protein of interest. Proteins were incubated with silica beads for 20 min at RT on a plate shaker to avoid beads from settling to bottom of well. If proteins were already fluorescently-tagged with a known dye-protein ratio then the SLB-covered beads were resuspended in 200μl of flow cell buffer and run on a flow cytometry machine (LSII or FORTESSA) along with MESF standard beads, Quantum Alexa Fluor 488 MESF or Quantum Alexa Fluor 647 MESF (Bangs Laboratories). If unlabelled proteins were tested, silica beads were stained with saturating concentration of antibodies against the specific ligand reconstituted on the SLB-coated beads. Silica beads with no protein or DOPC-covered silica beads were prepared as controls. MESF standard beads were used to translate the mean fluorescence intensity of protein-coated SLB-silica beads to number of dye per bead. Then using the dye-protein ratio of the ligand tested, we calculated the mean number of ligands per beads. The surface area of a bead was known hence the density of ligands was calculated by dividing the mean number of ligands per bead by the surface area of the bead. If ligands were detected with anti-ligand mAb, then, we used either the dye-protein ratio of the mAb and assumed a 1:2 ratio for mAb:ligand to determine the mean number of ligand per bead. If the dye-protein ratio of antibody was not known, the Quantum Simply Cellular anti-mouse IgG kit was used to make a calibration standard curve (Bangs Laboratories, 815).

We found that the Quantum Simply Cellular anti-mouse IgG kit overestimates the antibody binding capacity of the included beads for monoclonal antibodies perhaps due to isotype specific binding by the capture antibodies and hence this resulted in systematic underestimation of specific activity. After communication with the supplier, we were informed the Quantum Simply Cellular anti-mouse IgG kit beads were calibrated using polyclonal mouse IgG. The supplier assumes that monoclonal antibodies of any isotype would be bind with equal capacity to Quantum Simply Cellular anti-mouse IgG beads; but the supplier didn’t confirm this experimentally. After some inconsistent results we decided to experimentally test this. We cross-tested the AF647 MESF beads against the Quantum Simply Cellular anti-mouse IgG kit from the same supplier (Bangs) using samples of directly conjugated Alexa Fluor 647 conjugated mAb of different isotypes with spectroscopically determined dye to protein ratios. The MESF beads were used to generate a standard curve for number of dye molecules and the corresponding mean fluorescent intensity they produce on a flow cytometer. This allowed us to determine the mean number of dye molecules on each bead population of the Quantum Simply Cellular anti-mouse IgG kit. Knowing the mean number of dye molecules on each bead population and the dye to protein ratio of each mAb allowed us to calculate the number of antibodies captureach by each Quantum Simply Cellular anti-mouse IgG bead population directly for different IgG isotypes under saturating conditions. We determined that the binding capacity of the Quantum Simply Cellular anti-mouse IgG kit was overestimated by an average of 5.6-fold for any given mouse IgG isotype and we applied the appropriate correction depending upon the isotype to our unknown conjugated mAb for absolute quantification. Given that this is a large correction that will likely impact biological conclusions, we suggest that investigators who must use the Quantum Simply Cellular anti-mouse IgG kit, or similar kits, for absolute quantification perform their own testing for the capacity for relevant isotypes, using this traceable approach. We have previously verified by comparison with binding of radioiodinated antibodies that the 488 and 647 MESF beads are accurate standards.

### Processing of CRC samples

CRC tissue was collected from colorectal cancer patients that underwent colectomy at the Oxford University Hospitals NHS Foundation Trust. The samples were collected under the Ethics 11/YH/0020 and 16/YH/0247 for the BRC Oxford GI Biobank - supported by NIHR Biomedical Research Center, Oxford.

Tumour tissue was cut into small pieces and digested with Collagenase A, 1 mg/ml (Sigma Aldrich) prepared in ADF+++ (DMEM/F12, 200mM GLutamax, 1M HEPES and 1× PenStrep) on a shaker at 37°C, shaking for 1 hr. At the end of 1hr incubation, the digested tissue was passed through a 70μm mesh filter sequentially to generate a single cell suspension. Cells were washed in 1XHBSS/1%PenStrep. Red blood cells lysis was performed and the cells were counted for downstream surface staining and flow cytometric analysis.

### Image Analysis

TIRF image analysis was performed in Fiji (ImageJ, version 2.0.0) and analysis presented in GraphPad Prism 7. For quantification analysis, background signal was subtracted from IS image, by acquiring the same area of a nearby position void of cells. For Airyscan© confocal images initial processing was carried out in Zen© software (Zeiss), 3D rendering was performed in Imaris© software (Bitplane), and final panel preparation in Fiji© software (http://Fiji/sc).

## References

1. Sanchez-Madrid, F. et al. Three distinct antigens associated with human T-lymphocyte-mediated cytolysis: LFA-1, LFA-2, and LFA-3. Proceedings of the National Academy of Sciences of the United States of America 79, 7489–7493 (1982).

2. Sanders, M.E. et al. Human memory T lymphocytes express increased levels of three cell adhesion molecules (LFA-3, CD2, and LFA-1) and three other molecules (UCHL1, CDw29, and Pgp-1) and have enhanced IFN-gamma production. Journal of immunology (Baltimore, Md.: 1950) 140, 1401–1407 (1988).

3. Wang, J.H. et al. Structure of a heterophilic adhesion complex between the human CD2 and CD58 (LFA-3) counterreceptors. Cell 97, 791–803 (1999).

4. Bockenstedt, L.K. et al. The CD2 ligand LFA-3 activates T cells but depends on the expression and function of the antigen receptor. Journal of immunology (Baltimore, Md.: 1950) 141, 1904–1911 (1988).

5. Espagnolle, N. et al. CD2 and TCR synergize for the activation of phospholipase Cgamma1/calcium pathway at the immunological synapse. International immunology 19, 239–248 (2007).

6. Zaru, R., Cameron, T.O., Stern, L.J., Muller, S. & Valitutti, S. Cutting edge: TCR engagement and triggering in the absence of large-scale molecular segregation at the T cell-APC contact site. Journal of immunology (Baltimore, Md.: 1950) 168, 4287–4291 (2002).

7. Kaizuka, Y., Douglass, A.D., Vardhana, S., Dustin, M.L. & Vale, R.D. The coreceptor CD2 uses plasma membrane microdomains to transduce signals in T cells. The Journal of cell biology 185, 521–534 (2009).

8. van der Merwe, P.A. A subtle role for CD2 in T cell antigen recognition. The Journal of experimental medicine 190, 1371–1374 (1999).

9. Bachmann, M.F., Barner, M. & Kopf, M. CD2 sets quantitative thresholds in T cell activation. The Journal of experimental medicine 190, 1383–1392 (1999).

10. Carmo, A.M., Mason, D.W. & Beyers, A.D. Physical association of the cytoplasmic domain of CD2 with the tyrosine kinases p56lck and p59fyn. European journal of immunology 23, 2196–2201 (1993).

11. A role in transmembrane signaling for the cytoplasmic domain of the CD2 T lymphocyte surface antigen.

12. Skanland, S.S., Moltu, K., Berge, T., Aandahl, E.M. & Tasken, K. T-cell co-stimulation through the CD2 and CD28 co-receptors induces distinct signalling responses. The Biochemical journal 460, 399–410 (2014).

13. Pandolfi, F. et al. Expression of cell adhesion molecules in human melanoma cell lines and their role in cytotoxicity mediated by tumor-infiltrating lymphocytes. Cancer 69, 1165–1173 (1992).

14. Patel, S.J. et al. Identification of essential genes for cancer immunotherapy. Nature 548, 537–542 (2017).

15. McKinney, E.F., Lee, J.C., Jayne, D.R., Lyons, P.A. & Smith, K.G. T-cell exhaustion, co-stimulation and clinical outcome in autoimmunity and infection. Nature 523, 612–616 (2015).

16. De Jager, P.L. et al. The role of the CD58 locus in multiple sclerosis. Proceedings of the National Academy of Sciences of the United States of America 106, 5264–5269 (2009).

17. Raychaudhuri, S. et al. Genetic variants at CD28, PRDM1 and CD2/CD58 are associated with rheumatoid arthritis risk. Nature genetics 41, 1313–1318 (2009).

18. Abdul Razak, F.R., Diepstra, A., Visser, L. & van den Berg, A. CD58 mutations are common in Hodgkin lymphoma cell lines and loss of CD58 expression in tumor cells occurs in Hodgkin lymphoma patients who relapse. Genes and immunity 17, 363–366 (2016).

19. Hui, E. et al. T cell costimulatory receptor CD28 is a primary target for PD-1-mediated inhibition. Science (New York, N.Y.) 355, 1428–1433 (2017).

20. Yokosuka, T. et al. Programmed cell death 1 forms negative costimulatory microclusters that directly inhibit T cell receptor signaling by recruiting phosphatase SHP2. The Journal of experimental medicine 209, 1201–1217 (2012).

21. Zinselmeyer, B.H. et al. PD-1 promotes immune exhaustion by inducing antiviral T cell motility paralysis. The Journal of experimental medicine 210, 757–774 (2013).

22. Acuto, O. & Michel, F. CD28-mediated co-stimulation: a quantitative support for TCR signalling. Nature reviews. Immunology 3, 939–951 (2003).

23. Thompson, C.B. et al. CD28 activation pathway regulates the production of multiple T-cell-derived lymphokines/cytokines. Proceedings of the National Academy of Sciences of the United States of America 86, 1333–1337 (1989).

24. Miller, B.C. et al. Subsets of exhausted CD8(+) T cells differentially mediate tumor control and respond to checkpoint blockade. Nature immunology 20, 326–336 (2019).

25. Doering, T.A. et al. Network analysis reveals centrally connected genes and pathways involved in CD8+ T cell exhaustion versus memory. Immunity 37, 1130–1144 (2012).

26. Wherry, E.J. et al. Molecular signature of CD8+ T cell exhaustion during chronic viral infection. Immunity 27, 670–684 (2007).

27. Chirica, M. et al. Phenotypic analysis of T cells infiltrating colon cancers: Correlations with oncogenetic status. Oncoimmunology 4, e1016698 (2015).

28. Agata, Y. et al. Expression of the PD-1 antigen on the surface of stimulated mouse T and B lymphocytes. International immunology 8, 765–772 (1996).

29. Abu-Shah, E. et al. A tissue-like platform for studying engineered quiescent human T-cells’ interactions with dendritic cells. Biorxiv (2019). https://doi.org/10.1101/587386

30. Monks, C.R., Freiberg, B.A., Kupfer, H., Sciaky, N. & Kupfer, A. Three-dimensional segregation of supramolecular activation clusters in T cells. Nature 395, 82–86 (1998).

31. Huang, J. et al. CD28 plays a critical role in the segregation of PKC theta within the immunologic synapse. Proceedings of the National Academy of Sciences of the United States of America 99, 9369–9373 (2002).

32. Davis, S.J. & van der Merwe, P.A. The kinetic-segregation model: TCR triggering and beyond. Nature immunology 7, 803–809 (2006).

33. Ilani, T., Vasiliver-Shamis, G., Vardhana, S., Bretscher, A. & Dustin, M.L. T cell antigen receptor signaling and immunological synapse stability require myosin IIA. Nature immunology 10, 531–539 (2009).

34. Papa, I. et al. TFH-derived dopamine accelerates productive synapses in germinal centres. Nature 547, 318–323 (2017).

35. Varma, R., Campi, G., Yokosuka, T., Saito, T. & Dustin, M.L. T cell receptor-proximal signals are sustained in peripheral microclusters and terminated in the central supramolecular activation cluster. Immunity 25, 117–127 (2006).

36. Gruber, I.V. et al. Down-regulation of CD28, TCR-zeta (zeta) and up-regulation of FAS in peripheral cytotoxic T-cells of primary breast cancer patients. Anticancer research 28, 779–784 (2008).

37. Yokosuka, T. et al. Spatiotemporal regulation of T cell costimulation by TCR-CD28 microclusters and protein kinase C theta translocation. Immunity 29, 589–601 (2008).

38. Green, J.M., Karpitskiy, V., Kimzey, S.L. & Shaw, A.S. Coordinate regulation of T cell activation by CD2 and CD28. Journal of immunology (Baltimore, Md.: 1950) 164, 3591–3595 (2000).

39. Milstein, O. et al. Nanoscale increases in CD2-CD48-mediated intermembrane spacing decrease adhesion and reorganize the immunological synapse. The Journal of biological chemistry 283, 34414–34422 (2008).

40. Campi, G., Varma, R. & Dustin, M.L. Actin and agonist MHC-peptide complex-dependent T cell receptor microclusters as scaffolds for signaling. The Journal of experimental medicine 202, 1031–1036 (2005).

41. Yokosuka, T. et al. Newly generated T cell receptor microclusters initiate and sustain T cell activation by recruitment of Zap70 and SLP-76. Nature immunology 6, 1253–1262 (2005).

42. Lamason, R.L., McCully, R.R., Lew, S.M. & Pomerantz, J.L. Oncogenic CARD11 mutations induce hyperactive signaling by disrupting autoinhibition by the PKC-responsive inhibitory domain. Biochemistry 49, 8240–8250 (2010).

43. Bunnell, S.C. et al. T cell receptor ligation induces the formation of dynamically regulated signaling assemblies. The Journal of cell biology 158, 1263–1275 (2002).

44. Choudhuri, K. et al. Polarized release of T-cell-receptor-enriched microvesicles at the immunological synapse. Nature 507, 118–123 (2014).

45. Shen, K., Thomas, V.K., Dustin, M.L. & Kam, L.C. Micropatterning of costimulatory ligands enhances CD4+ T cell function. Proceedings of the National Academy of Sciences of the United States of America 105, 7791–7796 (2008).

46. Davis, S.J. & van der Merwe, P.A. The structure and ligand interactions of CD2: implications for T-cell function. Immunology today 17, 177–187 (1996).

47. Kumari, S. et al. T Lymphocyte Myosin IIA is Required for Maturation of the Immunological Synapse. Frontiers in Immunology 3 (2012).

48. Hahn, W.C., Rosenstein, Y., Calvo, V., Burakoff, S.J. & Bierer, B.E. A distinct cytoplasmic domain of CD2 regulates ligand avidity and T-cell responsiveness to antigen. Proceedings of the National Academy of Sciences of the United States of America 89, 7179–7183 (1992).

49. Kivens, W.J. et al. Identification of a proline-rich sequence in the CD2 cytoplasmic domain critical for regulation of integrin-mediated adhesion and activation of phosphoinositide 3-kinase. Molecular and cellular biology 18, 5291–5307 (1998).

50. Hahn, W.C. & Bierer, B.E. Separable portions of the CD2 cytoplasmic domain involved in signaling and ligand avidity regulation. The Journal of experimental medicine 178, 1831–1836 (1993).

51. Hutchings, N.J., Clarkson, N., Chalkley, R., Barclay, A.N. & Brown, M.H. Linking the T cell surface protein CD2 to the actin-capping protein CAPZ via CMS and CIN85. The Journal of biological chemistry 278, 22396–22403 (2003).

52. Sharpe, A.H., Wherry, E.J., Ahmed, R. & Freeman, G.J. The function of programmed cell death 1 and its ligands in regulating autoimmunity and infection. Nature immunology 8, 239–245 (2007).

53. Wei, F. et al. Strength of PD-1 signaling differentially affects T-cell effector functions. Proceedings of the National Academy of Sciences of the United States of America 110, E2480–2489 (2013).

54. Lo, D.J. et al. Selective targeting of human alloresponsive CD8+ effector memory T cells based on CD2 expression. American journal of transplantation: official journal of the American Society of Transplantation and the American Society of Transplant Surgeons 11, 22–33 (2011).

55. Leitner, J., Herndler-Brandstetter, D., Zlabinger, G.J., Grubeck-Loebenstein, B. & Steinberger, P. CD58/CD2 Is the Primary Costimulatory Pathway in Human CD28-CD8+ T Cells. Journal of immunology (Baltimore, Md.: 1950) 195, 477–487 (2015).

56. Yao, X. et al. Isolation and Characterization of an HLA-DPB1*04: 01-restricted MAGE-A3 T-Cell Receptor for Cancer Immunotherapy. Journal of immunotherapy (Hagerstown, Md.: 1997) 39, 191–201 (2016).

57. Li, Y. et al. Directed evolution of human T-cell receptors with picomolar affinities by phage display. Nature biotechnology 23, 349–354 (2005).

58. Dustin, M.L., Starr, T., Varma, R. & Thomas, V.K. Supported planar bilayers for study of the immunological synapse. Current protocols in immunology Chapter 18, Unit 18.13 (2007).

59. Dustin, M.L. et al. Quantification and modeling of tripartite CD2-, CD58FC chimera (alefacept)-, and CD16-mediated cell adhesion. The Journal of biological chemistry 282, 34748–34757 (2007).

