## Supplemental Information for "CD2 expression acts as a quantitative checkpoint for immunological synapse structure and T-cell activation"

### SUPPLEMENTARY INFORMATION

**Supplementary Figure 1. CD8 exhaustion signature, CD2 gene expression and levels in $CD4^+$ and $CD8^+$ CRC TILs**

The putative negative correlation of T cell exhaustion genesets and CD2 expression described in Figure 1 was then validated using the METALLIC colorectal cancer clinical cohort **a)** The correlation between exhausted $CD8^+$ T-cell gene signature and CD2 expression is shown for (top) "exhausted vs effector $CD8^+$ T cell gene signature" and (bottom) "exhausted vs naive $CD8^+$ T cell gene signature". **b)** The mean number of CD2 molecules/cell, for $CD4^+$ TIL subsets from Figure 1(b-c), is shown for each patient. Patients were sorted from highest to lowest CD2 expression of the $CD127^-PD\text{-}1^+$ $CD8^+$ T-cell compartment. **c)** Comparison of CD2 levels (mean±S.D.) in $CD127^-PD\text{-}1^+$ $CD8^+$ T-cells of Group A ($4.8x10^4 \pm 7.6x10^3$), 14 patients (from Figure 1, showing the $CD2^{low}$ phenotype) and Group B ($9.0x10^4 \pm 1.5x10^4$), 5 patients CRC with CD2 levels detected in peripheral blood $CD8^+$ T-cell of healthy individuals (naïve, $4.8x10^4 \pm 8.9x10^3$; memory, $7.6x10^4 \pm 8.9x10^3$). ****, $p<0.0001$; ***, $p<0.006$ with non-parametric Mann Whitney *U*-test. Error bars represent SEM. **d)** CD2 against PD-1 levels of $CD127^-PD\text{-}1^+$ $CD4^+$ TILs from (b) are shown for each donor.

Dotted line crossing x-axis (with grey rectangle) represents the average PD-1 levels (±SD) expressed in memory $PD\text{-}1^+$ $CD3^+CD4^+$ T cells found in peripheral blood of healthy controls. Dashed or dotted line crossing y-axis (with grey rectangle) represents the average CD2 levels (±SD) expressed in naive and memory $CD3^+CD4^+$ T cells, respectively, found in peripheral blood of healthy individuals.

**Supplementary Figure 2. A unique ring pattern, “corolla”, formed by CD2-CD58 interactions in the IS.**

**a)** Further examples of the CD2 and LFA-1 signal in fixed T:B conjugates as in Figure 1. Shown in 3D rendering (IMARIS software), are T:B cell conjugates (top panels) and 1 μm thick slice of the corresponding T:B cell interface shown in the bottom panels. Images were captured on an Airy-Scan Confocal Microscope (ZEISS). Representative images are shown from two independent experiments. Scale bar, 5 μm. **b)** Representative histograms showing the CD58 expression levels of the human CF996 EBV-transformed B cells (solid empty), human peripheral blood monocytes (light grey), *in vitro* differentiated immature monocyte-derived dendritic cells (dark grey) and *in vitro* maturate monocyte-derived dendritic cells (black). Dotted histogram represents isotype control stained cells. Scale bar, 5 μm.

**Supplementary Figure 3. pMHC-induced CD2 corolla captures ligated CD28.**

*1G4*$^{+}$ TCR CD8$^{+}$ T cells incubated on ICAM-1 (200/μm$^{2}$), NY-ESO-9V-peptide-loaded HLA-A2 (30/μm$^{2}$), CD80 (200/μm$^{2}$) with CD58 (200/μm$^{2}$) reconstituted SLB and real-time imaged with TIRFM, 10-15 min after contact and/or fixed at 15 min of incubation. Fluorescently labelled streptavidin was used to track the biotinylated NY-ESO-9V-peptide-loaded MHC. A representative image from two independent experiments is shown. Scale bar, 5 μm.

**Supplementary Figure 4. CD2 expression determines corolla formation.**

**a)** The gating strategy, in PBMCs, to quantify the levels of surface CD2 in T cell subsets from healthy individuals is shown. After gating on single and live cells, CD4$^{+}$CD3$^{+}$ (left) or CD8$^{+}$CD3$^{+}$ (right) cells were selected and divided into naïve

($CD62L^+CD45RA^+$), central memory ($CD62L^+CD45RA^-$), effector memory ($CD62L^-CD45RA^-$) and effector memory that re-expressed CD45RA ($CD62L^-CD45RA^+$). **b)** The histograms show an example of AND T cells transfected with different levels of human CD2 (hCD2FL; full length protein) and compared to CD2 levels found in human peripheral blood T cells from healthy individuals. Control cells were stained untransfected AND T cells (empty solid line histogram). **c)** Transfected AND T cells from (b) after a 15 min incubation on ICAM-1 ($200/\mu m^2$), MCC-I-$E^k$ ($30/\mu m^2$) and CD58 ($200/\mu m^2$) reconstituted SLBs, fixed and imaged with TIRFM. The ICAM-1, CD2 and CD58 signals in the IS of a random selection of AND T cells are shown for one representative of two such experiments. **d)** Same as in (c) but using AND T cells transfected with hCD2TM (hCD2 lacking its cytoplasmic tail)

**Supplementary Figure 5. $CD127^-$PD-$1^+$ $CD8^+$ TILs from CRC patients are enriched in $CD28^-$ TILs while $CD127^-$PD-$1^+$ $CD4^+$ TILs are mainly $CD28^+$.**

**a)** (Left) Representative CD28 and PD-1 expression plots (and gating) in $CD127^-$PD-$1^+$ $CD4^+$ from one CRC patient. (Right) The proportion of CD28 positive ($CD28^+$) and CD28 negative ($CD28^-$) T cells present within the viable $CD127^-$PD-$1^+$ $CD4^+$ TILs is shown for each CRC patient. **b)** (Left) Representative CD28 and PD-1 expression plots (and gating) in $CD127^-$PD-$1^+$ $CD8^+$ from one CRC patient. (Right) The proportion of CD28 positive ($CD28^+$) and CD28 negative ($CD28^-$) T cells present within the viable $CD127^-$PD-$1^+$ $CD8^+$ TILs is shown for each CRC patient.

**Supplementary Movie 1 and 2 (connects to Figure 2)**

Tracking of IS formation by a human T-cell incubated on ICAM1 (200/$\mu m^2$), anti-CD3 Fab (30/$\mu m^2$), CD58 (200/$\mu m^2$) reconstituted SLBs. Cells were imaged at 4s intervals with TIRFM. Scale bar

**Supplementary Movie 3 (connects to Figure 3)**

Tracking of IS formation by a human T-cell incubated on ICAM1 (200/$\mu m^2$), anti-CD3 Fab (30/$\mu m^2$), CD58 (200/$\mu m^2$), CD80 (100/$\mu m^2$) reconstituted SLBs. Cells were imaged at 4s intervals with TIRFM.

**Supplementary Table 1a. Clinical characteristics of CRC patients assessed in this study.**

| Patient ID | Type | Age | Gender | Specific Stage of disease | Tumour side (left or right side? | Is it colon or rectal? | Prognosis/recurrence? |
|---|---|---|---|---|---|---|---|
| 1 | CRC | 84 | M | T4 N1 M0 | Right | Ascending colon | pT3 pN0 Ly0 V0 R0. No recurrent or metastatic disease is identified. |
| 2 | CRC | 70 | F | T3 N0 M0 | Left | Sigmoid colon | pT3 pN2 Ly0 V0 R0. No evidence of residual or recurrent disease. |
| 3 | CRC | 75 | F | T2N0 | Left | Rectosigmoid colon | ypT3 ypN1 Ly0 V1 R0. No evidence of local or distant tumour or nodal recurrence. |
| 4 | CRC | 65 | M | T3 N0 M0 | ND | Rectum below perit.reflexion | pT3 pN2 Ly1 V1 R1. Further interval disease progression in multiple lung metastases. |
| 5 | CRC | 66 | M | T3 N2 M0 | Right | Caecal cancer | pT3 pN0 Ly0 V0 R0. No evidence of disease recurrence. |
| 6 | CRC | 71 | M | Tx | Right | Transverse colon | pT3 pN0 Ly0 V0 R0. No evidence of disease recurrence |
| 7 | CRC | 39 | M | T2 N0 M0 | Left | Rectosigmoid colon | pT3 pN0 Ly0 V0 R0. No metastatic disease. |
| 8 | CRC | 75 | F | T2 N0 M0 | ND | Rectum below perit.reflexion | pT2 pN0 Ly1 V0. |
| 9 | CRC | 84 | M | T4 N1 M0 | Left | Sigmoid colon | pT4b pN1a Ly1 V1 R0. |
| 10 | CRC | 68 | M | T2 N0 M0 | ND | Rectum below perit.reflexion | pT3 pN0 Ly0 V1 R0. New small bowel dilatation and mild small bowel thickening. |
| 11 | CRC | 55 | F | T3 N2 M0 | Left | Descending colon | ND |
| 12 | CRC | 76 | F | T4 | Right | Transverse colon | pT4b,N0,L0,V0,R0 |
| 13 | CRC | 82 | M | T2 N0 M0 | Right | Caecum | ND |
| 14 | CRC | 83 | F | T2 N0 M0 | Right | Ascending colon | ND |

ND, no data available

**Supplementary Table1a. ctd.**

| Patient ID | Type | Age | Gender | Specific Stage of disease | Tumour side (left or right side? | Is it colon or rectal? | Prognosis/recurrence? |
|---|---|---|---|---|---|---|---|
| 15 | CRC | 83 | M | T2 N0 M0 | Left | Sigmoid colon | ND |
| 16 | CRC | 76 | F | T2 N0 M0 | Right | Caecum | ND |
| 17 | CRC | 81 | M | T3 N1 M0 | Right | Ascending colon | ND |
| 18 | CRC | 74 | F | T3 N1 Mx | Right | Hepatic flexure tumour | ND |
| 19 | CRC | 81 | F | T2/3a N1 V0 M0. | ND | Rectum below perit.reflexion | ND |

ND, no data available

### Supplementary Table 1b. Clinical characteristics of CRC patients assessed in this study.

| Patient ID | Genetic information |
|---|---|
| 1 | Mutation detected in the KRAS gene (c.38G>A, p.(Gly13Asp), COSM532). Immunohistochemistry for Mismatch repair proteins show no loss of nuclear staining (MLH1, MSH2, MSH4 and PMS1) |
| 2 | Mutation detected in the KRAS gene (c.35G>T, p.(Gly12Val), COSM520). Mutation detected in the TP53 gene (c.267_267delC, p.(Ser90fs), COSM1268330). |
| 3 | Mutation not detected in the KRAS, NRAS and BRAF genes. Mutations detected in the TP53 gene (c.841G>A, p.(Asp281Asn), COSM43596) and (c.817C>T, p. (Arg273Cys), COSM10659) |
| 4 | Mutation not detected in the KRAS and NRAS genes. |
| 5 | Mutation not detected in the KRAS, NRAS and BRAF genes. Mutation detected in the PIK3CA gene (c.3140A>G, p.(His1047Arg), COSM775). |
| 6 | Mutation detected in the KRAS gene (c.436G>A, p.(Ala146Thr), COSM19404). Mutations detected in the TP53 gene (c.328C>T, p.(Arg110Cys), COSM43682; c.916C>T, p.(Arg306Ter), COSM10663; and c.413C>T, p.(Ala138Val), COSM43818). Mutations detected in the PIK3CA gene (c.263G>A, p.(Arg88Gln), COSM746 and c.3140A>G, p. (His1047Arg), COSM775). |
| 7 | Mutation not detected in the KRAS, NRAS and BRAF genes. Mutation detected in the TP53 gene (c.524G>A, p.(Arg175His), COSM10648). |
| 8 | Mutation detected in the KRAS gene (c.35G>T, p.(Gly12Val), COSM520). Mutation not detected in the NRAS and BRAF genes. Mutations detected in the PIK3CA gene (c.1637A>G, p.(Gln546Arg), COSM12459). Mutation detected in the TP53 gene (c.742C>T, p.(Arg248Trp), COSM10656). |
| 9 | MMR immunohistochemistry shows intact expression. |
| 10 | Clinically significant variant not detected in the KRAS, NRAS or BRAF genes. Variant detected in the TP53 gene (c.832C>A, p.(Pro278Thr), COSM43697). |
| 11 | ND |
| 12 | Mutation not detected in the KRAS and NRAS genes. Mutation detected in the BRAF gene (c.1799T>A, p.(Val600Glu), COSM476). |
| 13 | Mutation detected in the KRAS gene (c.35G>A, p.(Gly12Asp), COSM521). |

ND, no data available

### Supplementary Table 1b. ctd.

| Patient ID | Genetic information |
|---|---|
| 14 | Mutation detected in the BRAF gene (c.1799T>A, p.(Val600Glu), COSM476). Presence of BRAF p.(Val600Glu) in the context of loss of MLH1 (IHC) is suggestive, although not conclusive, of a sporadic tumour (Molecular testing strategies for Lynch Syndrome in people with colorectal cancer. Mutation detected in the PIK3CA gene (c.263G>A, p.Arg88Gln), COSM746) |
| 15 | Mutation detected in the KRAS gene (c.35G>A, p.(Gly12Asp), COSM521). Mutation detected in the PIK3CA gene (c.1035T>A, p.(Asn345Lys), COSM754). |
| 16 | The tumour cells show positive immunostaining for MLH1, MSH2 and PMS2, but are immunonegative for MSH6, suggesting loss of expression. |
| 17 | MMR Immunohistochemistry:<br>MLH1: Absent<br>MSH2: Intact<br>MSH6: Absent<br>PMS2: Absent |
| 18 | The tumour cells are immunopositive for MSH2 and MSH6, but are negative for MLH1 and PMS2, suggesting loss of expression. |
| 19 | Mismatch repair immunostains show no loss of MLH1, PMS2, MSH2 and MSH6 NUCLEAR STAINING. |

**Supplementary Table 2. CD2 mean expression in natural T-cell subsets in peripheral blood from 12 healthy individuals.**

| T cell subset | Molecules/cell | (+/-)SD |
|---|---|---|
| Naïve CD4+ | 3.15E+04 | 4.44E+03 |
| CM CD4+ | 5.22E+04 | 7.39E+03 |
| EM CD4+ | 7.05E+04 | 1.23E+04 |
| CD45RA+ EM CD4+ | 4.02E+04 | 1.34E+04 |
| Naïve CD8+ | 4.79E+04 | 8.90E+03 |
| CM CD8+ | 7.46E+04 | 7.36E+03 |
| EM CD8+ | 7.79E+04 | 1.10E+04 |
| CD45RA+ EM | 5.62E+04 | 1.15E+04 |
| pooled CD45RA- (CM and EM) CD4+ | 6.13E+04 | 9.34E+03 |
| pooled CD45RA- (CM and EM) CD8+ | 7.63E+04 | 8.88E+03 |
| PD-1+ CD8+ (6 healthy individuals) | 7.78E+04 | 6.14E+03 |

CM, central memory; EM, effector memory.

**Supplementary Table 3. CD2 mean expression in CD127±PD-1± TIL subsets from 19 CRC patients.**

| T cell subset | Molecules/cell | (+/-)SD |
|---|---|---|
| $CD127^{+}PD\text{-}1^{-}\ CD4^{+}$ | 4.85E+04 | 1.27E+04 |
| $CD127^{+}PD\text{-}1^{+}\ CD4^{+}$ | 6.54E+04 | 1.50E+04 |
| $CD127^{-}PD\text{-}1^{+}\ CD4^{+}$ | 6.03E+04 | 1.59E+04 |
| $CD127^{-}PD\text{-}1^{-}\ CD4^{+}$ | 3.82E+04 | 7.10E+03 |
| $CD127^{+}PD\text{-}1^{-}\ CD8^{+}$ | 5.51E+04 | 1.17E+04 |
| $CD127^{+}PD\text{-}1^{+}\ CD8^{+}$ | 6.55E+04 | 1.86E+04 |
| $CD127^{-}PD\text{-}1^{+}\ CD8^{+}$ | 5.96E+04 | 2.11E+04 |
| $CD127^{-}PD\text{-}1^{-}\ CD8^{+}$ | 4.54E+04 | 1.04E+04 |

**Supplementary Table 4. PD-1 mean expression in natural T-cell subsets in peripheral blood from 12 healthy individuals.**

| T cell subset | Molecules/cell | (+/-)SD |
|---|---|---|
| CM CD4+ | 4.84E+03 | 1.57E+03 |
| EM CD4+ | 5.77E+03 | 1.52E+03 |
| CM CD8+ | 5.48E+03 | 1.28E+03 |
| EM CD8+ | 6.36E+03 | 1.26E+03 |
| pooled CD45RA- (CM and EM) CD4+ | 5.30E+03 | 1.53E+03 |
| pooled CD45RA- (CM and EM) CD8+ | 5.92E+03 | 1.23E+03 |

CM, central memory; EM, effector memory.

**Supplementary Table 5. PD-1 mean expression in $CD127^-PD-1^+$ TIL subsets from 19 CRC patients.**

| T cell subset | Molecules/cell | (+/-)SD |
|---|---|---|
| $CD127^-PD-1^+ CD4^+$ | 1.36E+04 | 6.34E+03 |
| $CD127^-PD-1^+ CD8^+$ | 1.45E+04 | 5.58E+03 |

**Supplementary Table 6. CD58 mean expression in a human cell line and human primary myeloid cells from peripheral blood of two healthy individuals.**

| **Cell type** | **CD58 molecules/cell** | (+/-)SD |
|---|---|---|
| CF996 B cells line | 8.98E+04 | ND |
| Monocytes | 5.30E+04 | 1.60E+03 |
| Monocyted-derived DCs (immature) | 6.33E+04 | 8.10E+03 |
| Monocyted-derived DCs (mature) | 1.15E+05 | 1.10E+04 |

ND, no data available

Fig. S1. CD8 exhaustion signature, CD2 gene expression and levels in CD4⁺ and CD8⁺ CRC TILs.

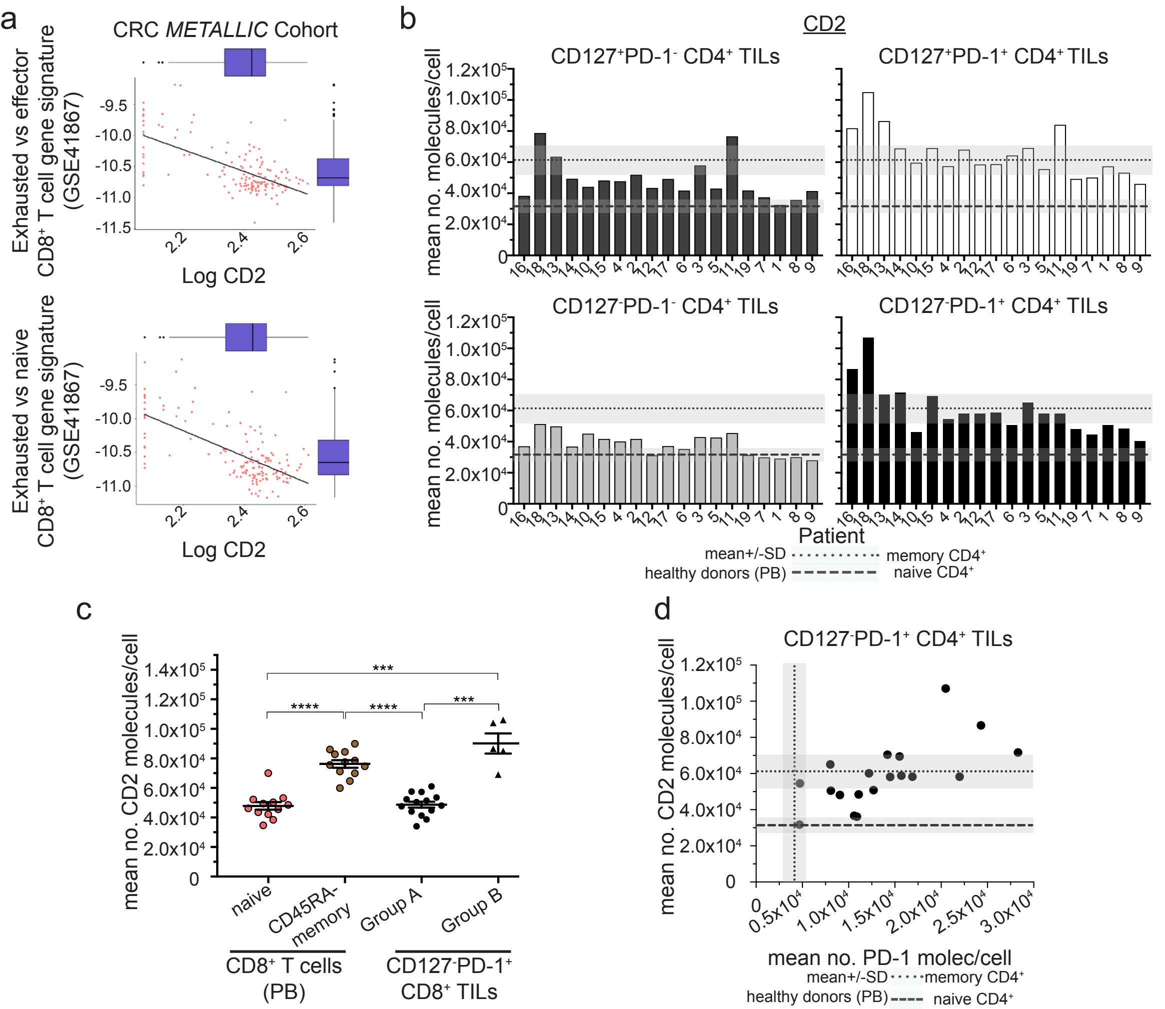

Fig. S2. A unique ring pattern, “corolla”, formed by CD2-CD58 interactions in the IS.

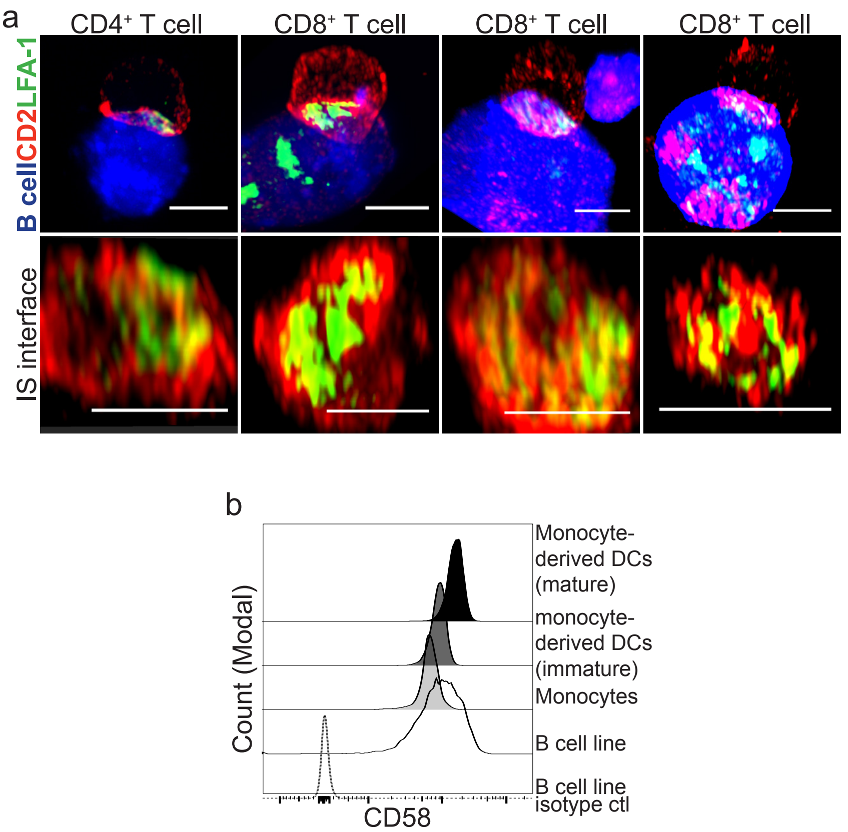

Fig. S3. pMHC-induced CD2 corolla captures ligated CD28

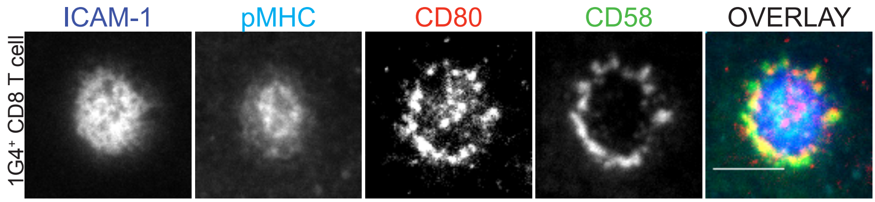

Fig. S4. CD2 expression levels determine corolla formation.

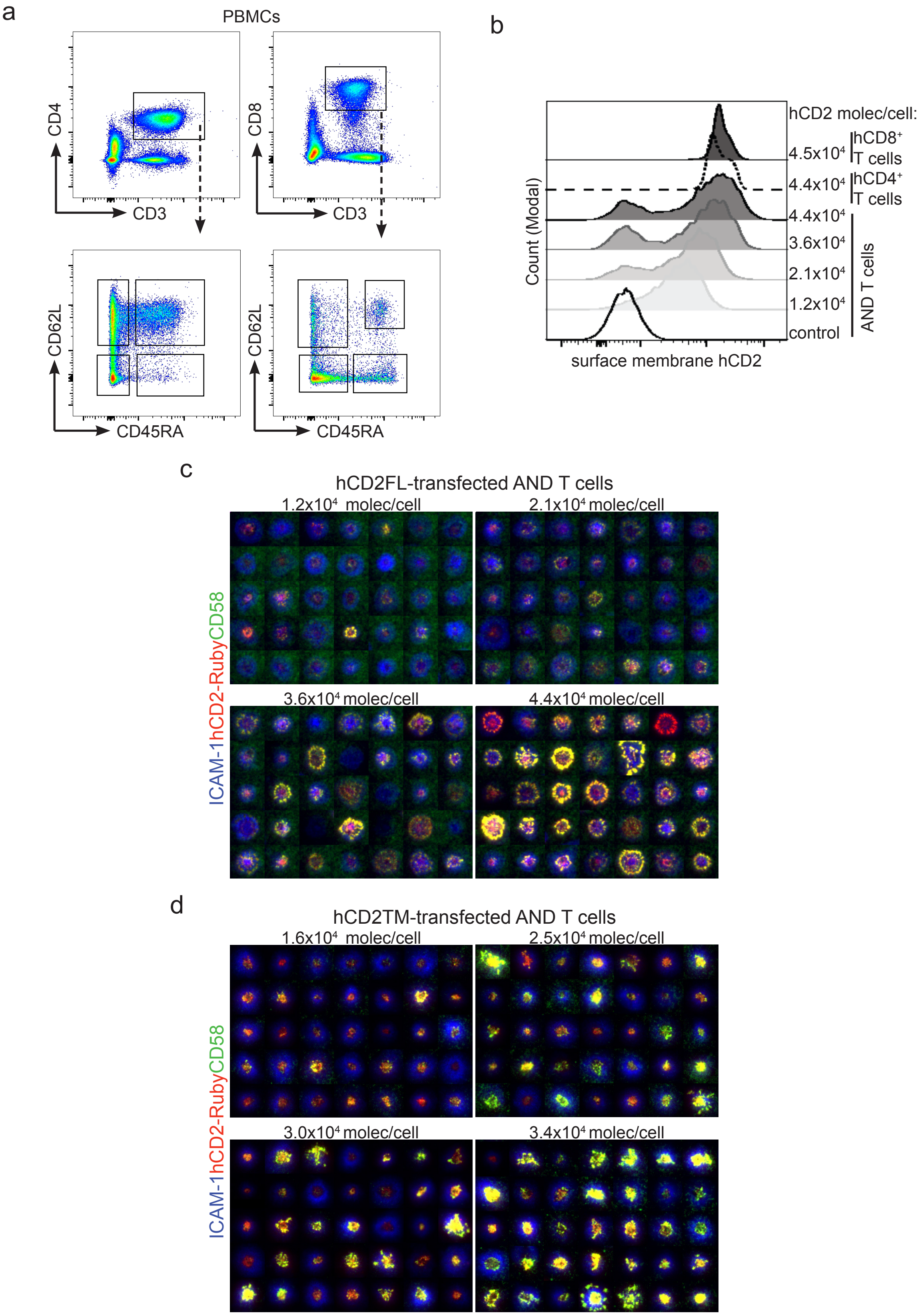

Fig. S5. $CD127^{-}PD\text{-}1^{+}$ $CD8^{+}$ TILs from CRC patients are enriched in $CD28^{-}$ TILs while $CD127^{-}PD\text{-}1^{+}$ $CD4^{+}$ TILs are mainly $CD28^{+}$.

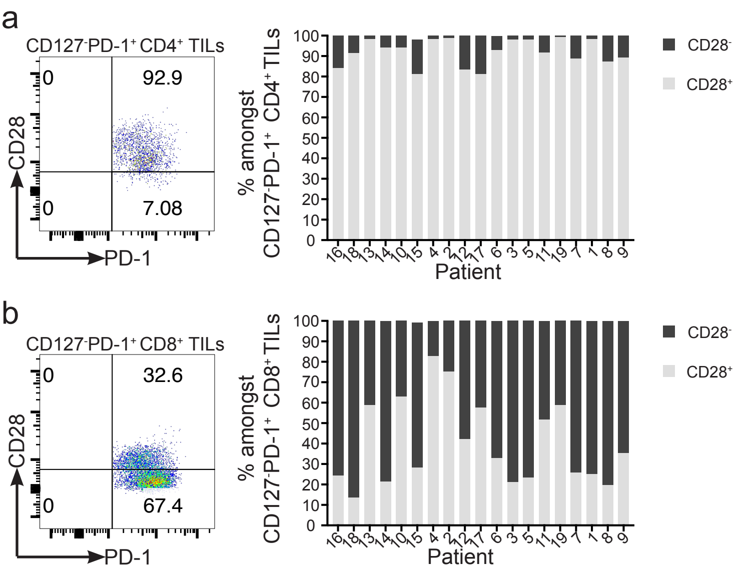
